# RNAfamProb Plus NeoFold: Estimations of Posterior Probabilities on RNA Structural Alignment and RNA Secondary Structures with Incorporating Homologous-RNA Sequences

**DOI:** 10.1101/812891

**Authors:** Masaki Tagashira, Kiyoshi Asai

## Abstract

**Motivation:** The *simultaneous optimization of the sequence alignment and secondary structures among RNAs, structural alignment*, has been required for the more appropriate comparison of functional *ncRNAs* than sequence alignment. *Pseudo-probabilities given RNA sequences on structural alignment* have been desired for more-accurate secondary structures, sequence alignments, consensus secondary structures, and structural alignments. However, any algorithms have not been proposed for these pseudo-probabilities.

**Results:** We invented the RNAfamProb algorithm, an algorithm for estimating these pseudo-probabilities. We performed the application of these pseudo-probabilities to two biological problems, the visualization with these pseudo-probabilities and *maximum-expected-accuracy secondary-structure* (estimation). The RNAfamProb program, an implementation of this algorithm, plus the NeoFold program, a maximum-expected-accuracy secondary-structure program with these pseudo-probabilities, demonstrated prediction accuracy better than three state-of-the-art programs of maximum-expected-accuracy secondary-structure while demanding running time far longer than these three programs as expected due to the intrinsic serious problem-complexity of structural alignment compared with independent secondary structure and sequence alignment. Both the RNAfamProb and NeoFold programs estimate matters more accurately with incorporating *homologous-RNA sequence*s.

**Availability:** The source code of each of these two programs is available on each of “https://github.com/heartsh/rnafamprob” and “https://github.com/heartsh/neofold”.

**Contact:** and.

**Supplementary information:** Supplementary data are available at *Bioinformatics* online.

## 1 Introduction

A large quantity of *RNAs not translated into proteins* (= *ncRNA*s) being functional have been discovered through sequencing technology including high-throughput sequencing technology (Maxam and Gilbert, 1977; Bentley *et al*., 2008). These RNAs are related to various biological processes such as epigenetic silencing (Pasmant *et al*., 2011), splicing regulation (Ji *et al*., 2003), translational control (Long and Caceres, 2009), apoptosis regulation, and cell cycle control (Kino *et al*., 2010). However, a large number of these RNAs are functionally unknown. RNAs fold into 3D structures while taking less *free energy* (= FE). Sets of *Base-Pairings* (= BPs) in these structures, *secondary structure*s (= SSs), should be considered together with RNA sequences when measuring the similarity of functional ncRNAs because both BP and *unpaired* bases play roles in biological processes (Wu *et al*., 1991).

The *optimization of residue alignments among biomolecules, sequence alignment* (Gotoh, 1982; Altschul *et al*., 1997; Katoh and Standley, 2013) (= SA), is required for the comparison of biomolecules mainly whose residues play roles in biological processes such as proteins and DNAs. The *simultaneous optimization of the SA and SSs among RNAs, STructural Alignment* (Sankoff, 1985) (= STA), has been required for the more appropriate comparison of functional ncRNAs. Pseudo-probabilities given RNA sequences on STA have been desired for more-accurate SSs, SAs, *consensus SS*s (= optimizations of aligning BP base-pairs among RNAs = CSSs) (Bernhart *et al*., 2008), and STAs. (Hamada *et al*., 2009c,a, 2011; Sato *et al*., 2012) However, any algorithms have not been proposed for estimating these pseudo-probabilities. Therefore, we invented the RNAfamProb algorithm, an algorithm for estimating pseudo-probabilities given RNA sequences on STA. We performed the application of these pseudo-probabilities to two biological problems, visualization with these pseudo-probabilities and *Maximum-Expected-Accuracy* (= MEA) *SS* (Hamada *et al*., 2009b) (estimation). The RNAfamProb program, an implementation of this algorithm, plus the NeoFold program, an MEA SS program with these pseudo-probabilities, demonstrated prediction accuracy better than three state-of-the-art MEA-SS-programs while demanding running time far longer than these three programs as expected due to the intrinsic serious problem-complexity of STA compared with independent SS and SA. Both the RNAfamProb and NeoFold programs estimate matters more accurately with incorporating *homologous-RNA sequence*s.

## 2 Methods

### 2.1 Our proposed method of maximum-expected-accuracy secondary structure

The workflow of our proposed MEA-SS-method, the NeoFold suite, is shown in Figure 1.

**Fig. 1.**
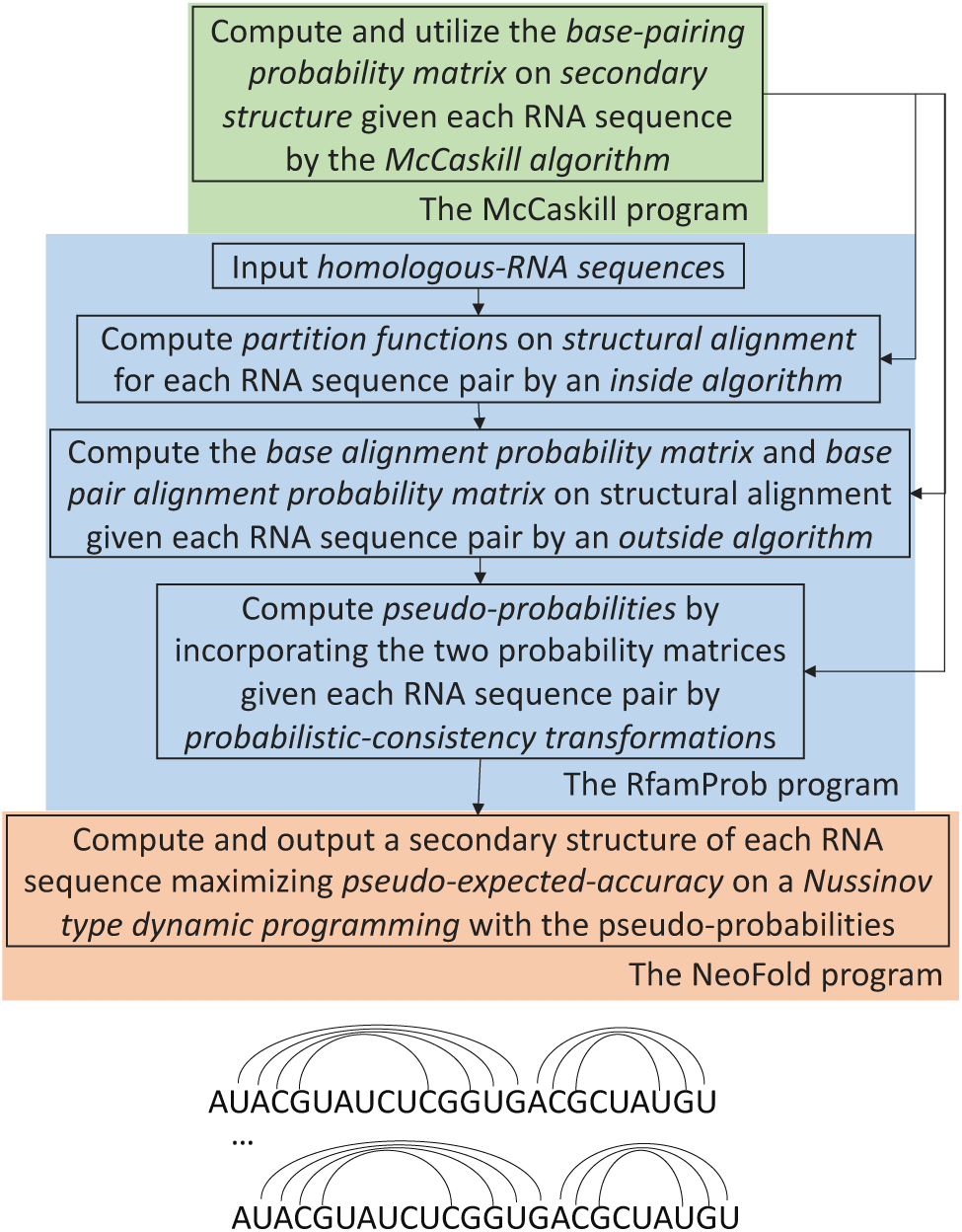
The workflow of our proposed method of maximum-expected-accuracy secondary-structure estimation, the Neofold suite. This suite combines the RNAfamProb, McCaskill, and NeoFold programs. The RNAfamProb program estimates pseudo-base-pairing-probabilities more accurate than base-pairing probabilities estimated by the McCaskill program by considering the structural alignment between two homologous-RNA sequences. The McCaskill program helps the RNAfamProb program finish computations in realistic time by filtering in possible base-pairings on an RNA sequence. The Neofold program estimates secondary structures more accurately by using these pseudo-probabilities instead of these probabilities.

### 2.2 Structural alignment

An STA between two RNA sequences ***RNA, RNA***′ is defined as 𝕊𝕋𝔸_ℝℕ𝔸:=[***RNA,RNA***′]_ := [***SS _RNA_, SS _RNA_***′, ***SA***_ℝℕ𝔸_] where ***SS***_***RNA***_ is defined as an SS of ***RNA*** and ***SA***_ℝℕ𝔸_ an SA of ℝℕ𝔸. The pseudo-positions 0, *N* + 1 are BP in any ***SS***_***RNA***_ where *N* is defined as the length of ***RNA***. Each of the pseudo-position pairs [0, 0], [*N* + 1, *M* + 1] is aligned in any ***SA***_ℝℕ𝔸_ where *M* is defined as the length of ***RNA***′. Two position pairs [*i, j*], [*k, l*]; *i, j, k, l ∈*N, 0 ≤ *i < j* ≤ *N* + 1, 0 ≤*k < l* ≤ *M* + 1 are *corresponding* if and only if each of [*i, j*], [*k, l*] are BP and [*i, j*], [*k, l*] are aligned. A position pair must be BP if this pair is aligned with a BP position pair. Only SSs without any *pseudoknot*s and *collinear* SAs are considered in this paper because considerations of these SSs and SAs lead to larger time and space complexities. (Sato *et al*., 2011; Bradley *et al*., 2009)

A pairwise CFG (= *pair-CFG*) describing STAs ℂ𝔽𝔾^sta^ is defined as

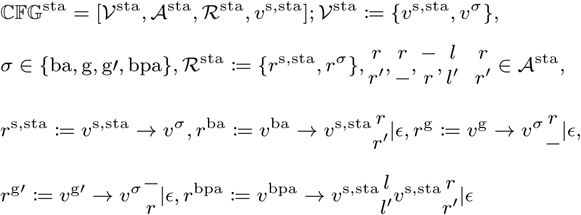

where each of *l, r, l′, r′* is defined as a base and −a gap. *𝒱*^sta^ is a set of *variable*s, *ℛ*^sta^ a set of transitions each from a variable to variables (= *rule*s), and *v*^s,sta^ a start variable. This pair-CFG is equivalent to the pair-CFG described in Dowell and Eddy (2006). An example of each of ***SS***_***RNA***_, ***SA***_ℝℕ𝔸_, 𝕊𝕋𝔸_ℝℕ𝔸_ is shown on each of (A), (B), and (C) in Figure 2.

**Fig. 2.**
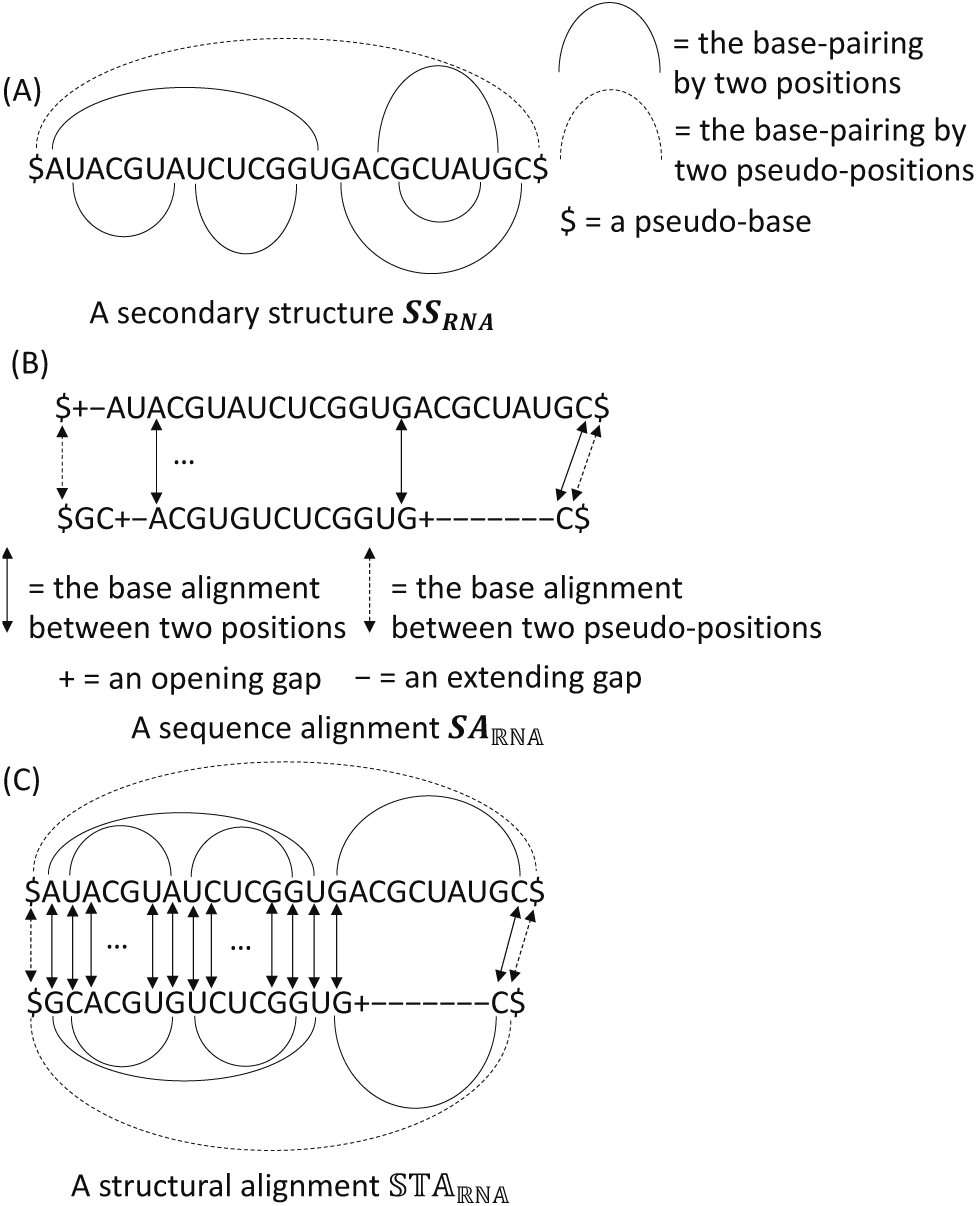
An example of each of (A) a secondary structure ***SS***_***RNA***_, (B) sequence alignment ***SA***_ℝℕ𝔸_, and (C) structural alignment STA_ℝℕ𝔸_.

### 2.3 Boltzmann distribution and posterior probabilities

𝔖𝔗𝔄_ℝℕ𝔸_ is defined as the *STA space* of ℝℕ𝔸. The Boltzmann distribution of 𝕊𝕋𝔸 *∈* 𝔖𝔗𝔄_RNA_ is defined as 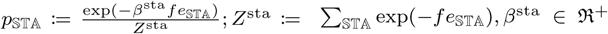 where *fe*_𝕊𝕋𝔸_ is defined as the FE of 𝕊𝕋𝔸 and ℜ ^+^ the positive-real-number space excluding 0. *Z*^sta^ is the *partition function* (= PF) on 𝔖𝔗𝔄_ℝℕ𝔸_ and *β*^sta^ a scale parameter of *fe*_𝕊𝕋𝔸_.

#### Definition 2.1.

*Each of the base alignment probability (= BAP) matrix (= BAPM) on STA and base pair alignment probability (= BPAP) matrix (= BPAPM) given* ℝℕ𝔸 *is defined as each of* 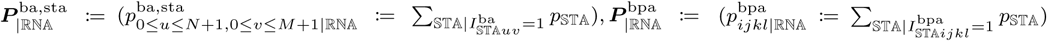 *where* 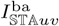 *is defined as 1 if the two positions u, v are aligned and unpaired in* 𝕊𝕋𝔸 *and 0 otherwise and* 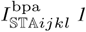 *if the two position pairs* [*i, j*], [*k, l*] *are corresponding in* 𝕊𝕋𝔸 *and 0 otherwise*.

### 2.4 Formulation of structural-alignment free energy *fe*_𝕊𝕋𝔸_

*fe*_𝕊𝕋𝔸_ can be formulated as

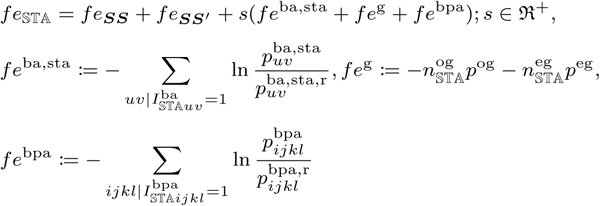

where *fe* _***SS***_ is defined as the FE of ***SS***, each of 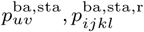 the *prior* BAP of the two positions *u, v* on STA on each of the two assumptions in whether ℝℕ𝔸 is correlated or uncorrelated, each of 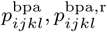 the prior BPAP of the two position pairs [*i, j*], [*k, l*] corresponding on each of these two assumptions, each of 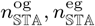 the number of each of opening and extending gaps in 𝕊𝕋𝔸, and each of *p*^*og*^, *p*^eg^ a penalty of each of two opening and extending gaps. *s* is the scale parameter of *fe*^ba,sta^ + *fe*^g^ + *fe*^bpa^ relative to *fe*_***SS***_ + *fe*_***SS*** ′_.

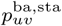 can be formulated as 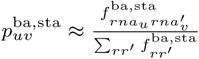 where 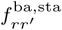 is defined as the frequency of *r, r ′* aligned in STAs for estimating 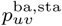 *rna*_*u*_ the *u*-th base of ***RNA***, and 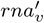 the *v*-th base of 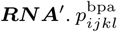 can be formulated similarly to 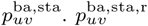 can be formulated as 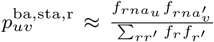 where *f*_*r*_ is defined as the frequency of *r* in STAs for estimating 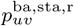.

Probabilities 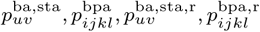 estimated are available as the RIBOSUM STA FE matrices. (Klein and Eddy, 2003) On the other hand, *fe*_***SS***_ is available as the parameters of the Turner model, a nearest-neighbor approximation model for RNA SS FE on thermodynamics. (Turner and Mathews, 2010) 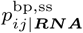 have been utilized as components of *fe*_***SS***_ on computing on STA due to computations simplified more though the suitability of 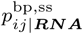 to *fe*_***SS***_ has not been discussed. (Hofacker *et al*., 2004; Do *et al*., 2008) In this study, the Turner model is combined with the two RIBOSUM STA FE 80-65 matrices for preventing the accuracy of 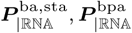 from losing.

### 2.5 *Inside-outside algorithm* for computing base alignment probability matrix and base pair alignment probability matrix 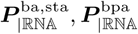

Each of 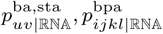 can be formulated as each of

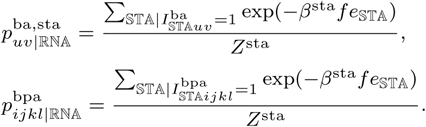

Thus, first *Z*^sta^ and then 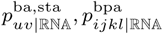 are formulated recursively, which leads to Algorithm 1.

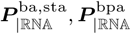 can be computed by Algorithm 1 with the time and space complexities *O*(*N*^4^ *M*^4^), *O*(*N*^2^ *M*^*2*^). This algorithm is the simultaneous solution of the Durbin (*forward-backward*) algorithm (Durbin *et al*., 1998) and McCaskill (inside-outside = IO) algorithm (McCaskill, 1990) and the IO algorithm version of the Sankoff algorithm, a maximum-likelihood (= ML) STA algorithm (Sankoff, 1985), as expected. An analysis of these two complexities is described in Supplementary section 2.

#### Algorithm 1 An inside-outside algorithm for computing a base alignment probability matrix and base pair alignment probability matrix 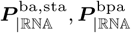.

1. **function** ioAlgo4BapmAndBpapm(ℝℕ𝔸)
2. Compute partition functions on an *inside algorithm* according to Supplementary section 1 // Inside step.
3. // Outside step. Partition functions other than partition functions 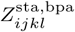, need to be recomputed for exploiting more sparsity of partition functions.
4. Compute base alignment probabilities and base pair alignment probabilities 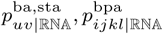 on an *outside algorithm* while recomputing partition functions other than 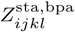 on this inside algorithm according to Supplementary section 2 and Supplementary section 1
5. **return** 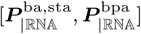

### 2.6 Inside-outside algorithm for computing base alignment probability matrix and base pair alignment probability matrix 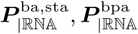 approximate

Algorithm 1 is not practical because of the time and space complexities of this algorithm *O*(*N* ^4^*M* ^4^), *O*(*N* ^2^*M* ^2^). IO algorithms for computing 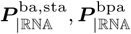 approximate (or sparse) with less time and space complexities can be gained by adding restrictions to (or sparsifying) *𝒮𝒯𝒜*_ℝℕ𝔸_.

Two restrictions frequently added to this space are the *minimum-BPP restriction* (Sato *et al*., 2012) and *maximum-gap-number restriction* (Torarinsson *et al*., 2007). 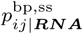 is defined as the BPP of the two positions given, *i, j* ***RNA*** on SS, computed by the McCaskill algorithm. The minimum-BPP restriction allows any two positions *i, j* to base-pair only when 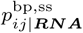 is greater than or equal to a minimum BPP *p*^bp,ss,m^ *∈* ℜ ^+^; 0 ≤ *p*^bp,ss,m^ ≤ 1. The maximum-gap-number restriction allows any STA only when the gap number in STA is less than or equal to a maximum gap-number *g*^m^ *∈* 𝔑; |*N* −*M* | ≤*g*^m^.

The probable-STA restriction is defined as the combination of the minimum-BPP and maximum-gap-number restrictions. 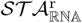 is defined as the STA space gained by adding the probable-STA restriction to any 𝕊𝕋𝔸. Each of 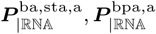 is defined as each of the approximate BAPM and BP APM gained by replacing *𝒮𝒯𝒜*_ℝℕ𝔸_ with 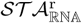 in Definition 2.2. Algorithm 1 with this restriction is virtually with the time and space complexities *O*(*L*^2^), *O*(*L*); *L* := max(*N, M*) if *p*^bp,ss,m^ takes a sufficiently large value and *g*^m^ takes a sufficiently small value.

### 2.7 Probabilistic-consistency transformation

A Probabilistic-Consistency Transformation (= PCT) is the transformation of a probability between a biomolucule or biomolecules and each homologous biomolecule into a pseudo-probability. (Do *et al*., 2005) This pseudo-probability contains information about all utilized homologous biomolecules. ***RF*** is defined as an *RNA family*, a sequence of homologous-RNA sequences each with a length of at most *N*. Each of 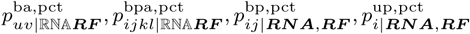 is defined as a pseudo-BAP, pseudo-BPAP, pseudo-BPP, and pseudo-unpairing-probability gained by performing a PCT between ℝℕ𝔸 or ***RNA*** and 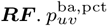 can be formulated as

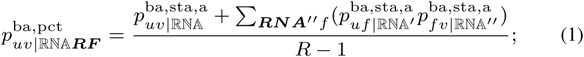

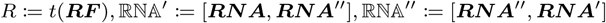

where ***RNA*** ″ ∈***RF*** (***RNA, RNA*** ′), 0 ≤*f* ≤ *t* (***RNA*** ″) −1 and *t*(***RF***) is defined as a function returning the type (or size) of ***RF***. 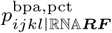 can be formulated similarly to 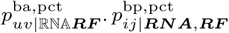 can be formulated as

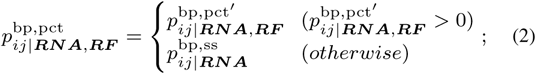

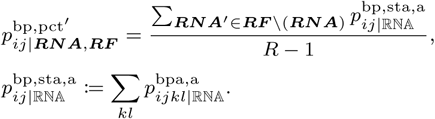

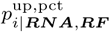 can be formulated as

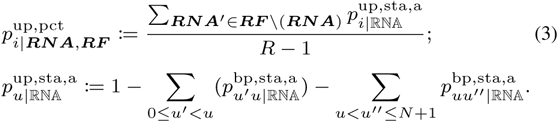

***RNA***_***RF***_ is defined as a sequence of pairs of the RNA sequences in ***RF***. Pseudo-probability matrices

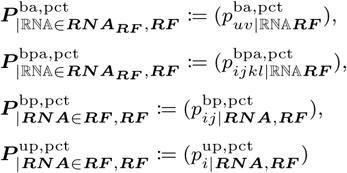

can be computed by the RNAfamProb algorithm, Algorithm 2.

#### Algorithm 2 The RNAfamProb algorithm.

1. **function** rnafamProb(***RF***)
2. Compute base alignment probability matrices and base pair alignment probability matrices 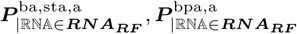 by Algorithm 1 with the probable-STA restriction
3. Compute pseudo-probabilities 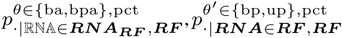 according to Equation 1, Equation 2, and Equation 3
4. **return** 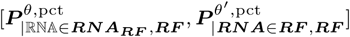

### 2.8 Maximum-expected-accuracy secondary structure incorporating single homologous-RNA sequence

The *accuracy* of ***SS*** against 𝕊 𝕋 𝔸 is defined as

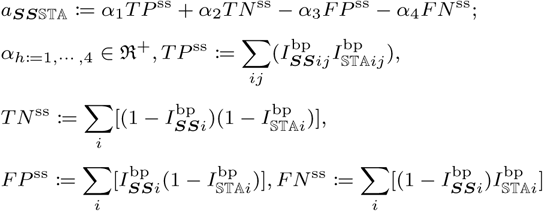

where 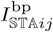 is defined as 1 if the two positions *i, j* are BP in 𝕊 𝕋 𝔸 and 0 otherwise.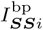 1 if *i* is BP in ***SS*** and 0 otherwise, and 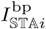 1 if *i* is BP in 𝕊 𝕋 𝔸 Each of *TP* ^ss^; *TN* ^ss^; *FP* ^ss^; *FN* ^ss^ is the number of each of true positives (= TPs), true negatives (= TNs), false positives (= FPs), and false negatives (= FNs). Each of *α*_*h*_ a scale parameter for each of *TP* ^ss^, *TN* ^ss^, −*FP* ^ss^, −*FN* ^ss^.

*a*_***SS***𝕊 𝕋 𝔸_ and accuracy 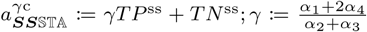 are equivalent in terms of SS accuracy. Expected accuracy 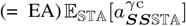 can be formulated as

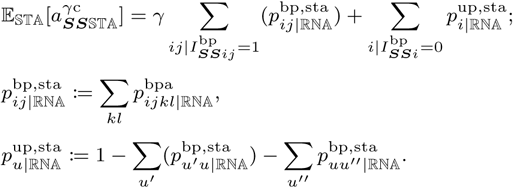

A proof of each of these relationship and formulation is described in each of Supplementary section 4 and Supplementary section 5.

#### Definition 2.2.

*The MEA SS of* ***RNA*** *is defined* as 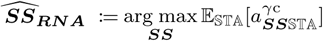.

*mea*_*ij*_ is defined as the EA of 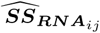 in the case in which the two positions *i, j* are BP where ***RNA***_*ij*_ is defined as the *substring* of ***RNA*** between *i, j* inclusive of *i, j*. max_***SS***_ 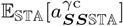. is gained from the fact max_***SS***_ 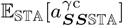.are equal for any ℝ ℕ 𝔸mea_*ij*_ can be formulated as

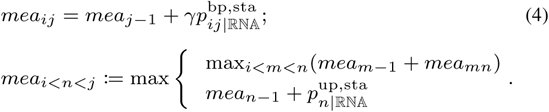

An interpretation of each of (A) the formulation of *mea*_*ij*_ and definition of *mean* is shown on each of (A) and (B) in Figure 3. It should be noted Equation 4 is a Nussinov type *dynamic programming* (= DP) recursive equation (Nussinov *et al*., 1978). *mea*_*i*_ is formulated (or initialized) as 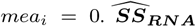 can be computed with the time and space complexities *O*(*N* ^4^), *O*(*N* ^2^) by Supplementary Algorithm 2. A proof of these two complexities is described in Supplementary section 6.

**Fig. 3.**
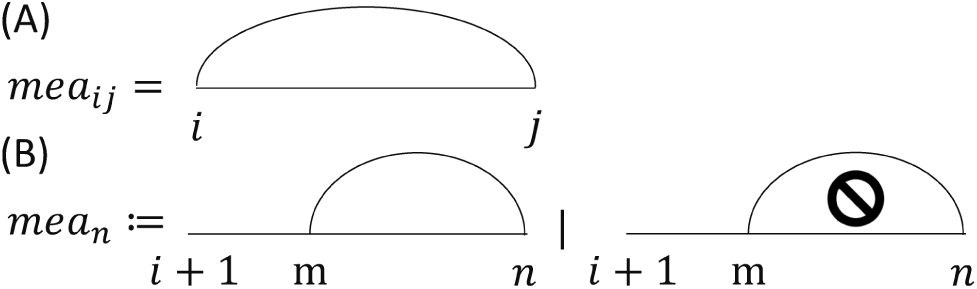
An interpretation of each of (A) the formulation of maximum expected accuracy *mea*_*ij*_ and (B) definition of maximum expected accuracy *mean*.

### 2.9 Pseudo-maximum-expected-accuracy secondary structure incorporating multiple homologous-RNA sequences

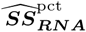 is defined as the MEA SS gained by replacing 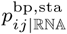 with 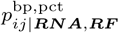 and 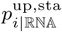with 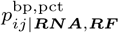 in Definition 2.2. 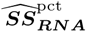 can be computed by the NeoFold algorithm, Supplementary algorithm 3. This algorithm is virtually with the time and space complexities *O*(*N* ^2^), *O*(*N*) if the number of positions able to base-pair with a position is sufficiently small.

### 2.10 Relationship between NeoFold algorithm and other maximum-expected-accuracy secondary-structure algorithms

The NeoFold algorithm can be considered as the CONTRAfold algorithm, an MEA SS algorithm (Do *et al*., 2006), improved by replacing probabilities on SS with pseudo-probabilities on STA and modifying slightly accuracy. This algorithm maximizes EA 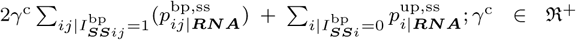. Thus, this algorithm counts twice TPs and FNs compared with the NeoFold algorithm. On the other hand, the NeoFold algorithm can be considered as the CentroidFold algorithm, an MEA SS algorithm (Hamada *et al*., 2009b), improved by replacing probabilities on SS with pseudo-probabilities on STA and modifying drastically accuracy. This algorithm maximizes EA

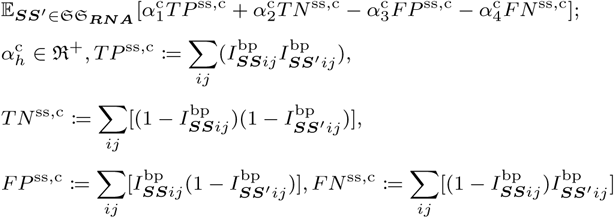

where 𝔖 𝔖_***RNA***_ is defined as the *SS space* of ***RNA***. This algorithm overcounts TNs compared with the NeoFold algorithm, especially when RNA sequences are long, since at most one position *j* can base-pair with a position *i*. The NeoFold algorithm counts TPs, TNs, FPs, and FNs equally, i.e. the contribution of these four numbers to the accuracy of this algorithm is unbiased.

The CentroidHomFold algorithm, an MEA SS algorithm incorporating homologous-RNA sequences via PCT similarly to the NeoFold algorithm, considers STAs by decomposing 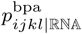 as 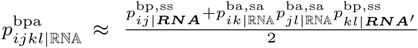 where 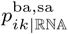 is defined as the BAP of the two positons *i, k* given ℝℕ𝔸 on SA, computed by the Durbin algorithm (Durbin *et al*., 1998). Mathematical decompositions like this decomposition are frequently utilized in many RNA informatics algorithms including the CentroidAlign algorithm (= an MEA SA algorithm considering STA) (Hamada *et al*., 2009a), CentroidAliFold algorithm (= an MEA CSS algorithm) (Hamada *et al*., 2011), and DAFS algorithm (= an MEA STA algorithm) (Sato *et al*., 2012) because precise considerations of STAs demand 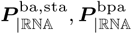, whose computations have been proposed in this paper. When the probable-STA restriction is removed completely on the RNAfamProb algorithm, the CentroidHomFold algorithm is an approximate version of the NeoFold algorithm.

The NeoFold algorithm incorporates precisely homologous-RNA sequences via STA whereas SS algorithms so far do not incorporate these sequences or avoid STA via mathematical-decomposition techniques as shown in Figure 4.

**Fig. 4.**
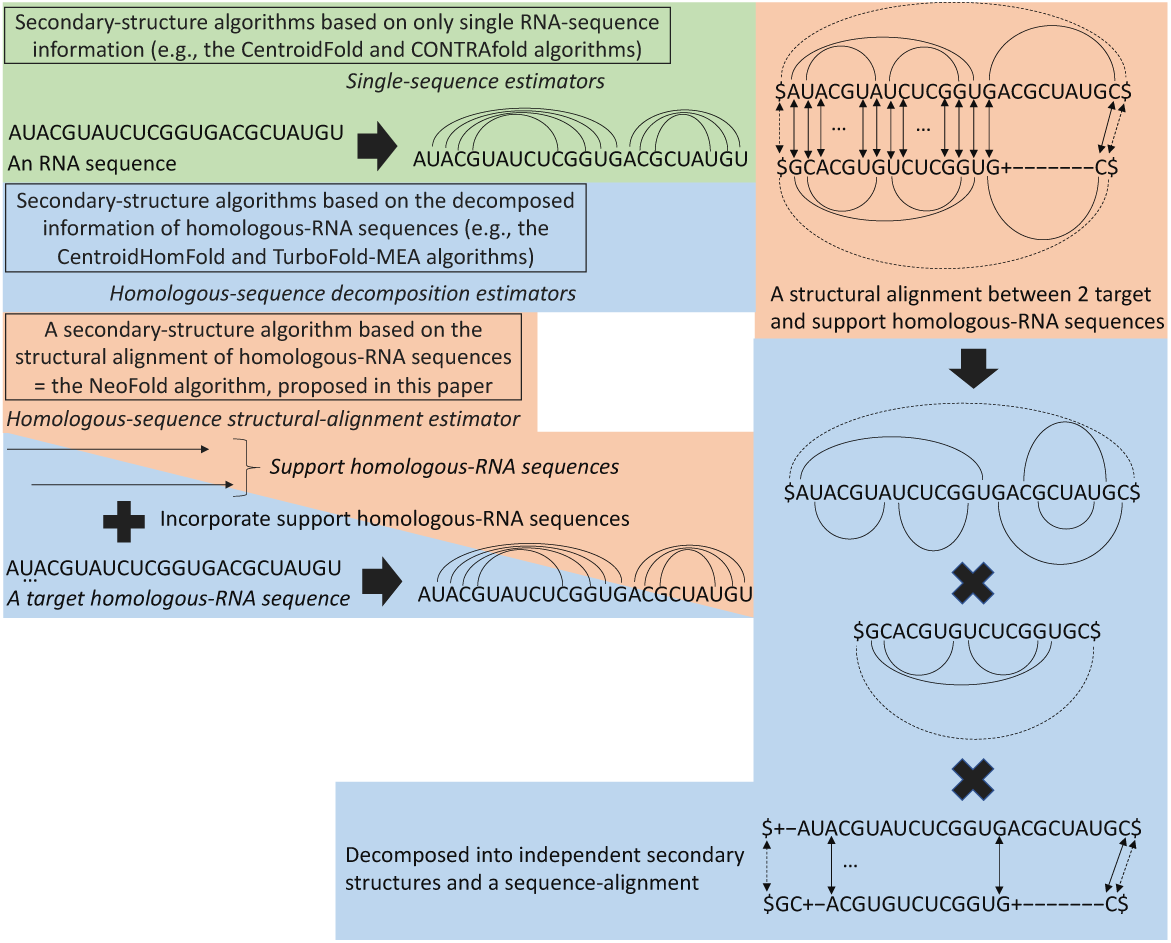
The comparison among three types of secondary-structure estimator. Single-sequence and homologous-sequence decomposition estimators are proposed in many studies so far. Homologous-sequence structural-alignment estimator has been proposed for the first time in this paper. The ascending order of the expected reliability of these three types is single sequence estimator → homologous-sequence decomposition estimator → homologous-sequence structural-alignment estimator.

## 3 Results and discussions

### 3.1 Algorithm implementations and environments for running programs

Each of the RNAfamProb, NeoFold, and McCaskill algorithms was implemented on the Rust programming language as the version 0.1.0 of each of the RNAfamProb, NeoFold, and McCaskill programs. These three programs employ multi-threading as much as possible for achieving running time as fast as possible. The source code of each of these three programs is available on each of “https://github.com/heartsh/rnafamprob”, “https://github.com/heartsh/neofold”, and “https://github.com/heartsh/rna-algos”. Programs were run on a computer composed of the “Intel Xeon CPU E5-2680 v2” CPU with 20 CPU logical cores and the clock rate 2.80 [GHz] and 64 [GB] of RAM unless environments for running programs are specified.

### 3.2 Visualization with pseudo-probabilities 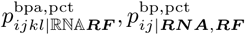

#### 3.2.1 Visualization of pseudo-base-pair-alignment-probability 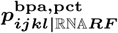 as reliability of two corresponding position-pairs [*i, j*], [*k,l*]

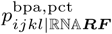 can be used as the reliability of the two corresponding position-pairs [*i, j*], [*k, l*] as in Figure 5. The ascending order of the expected reliability of the CSS on each of (A), (B), and (C) in Figure 5 is (A) → (B) → (C). This order is consistent with the observed reliability of these three CSSs.

**Fig. 5.**
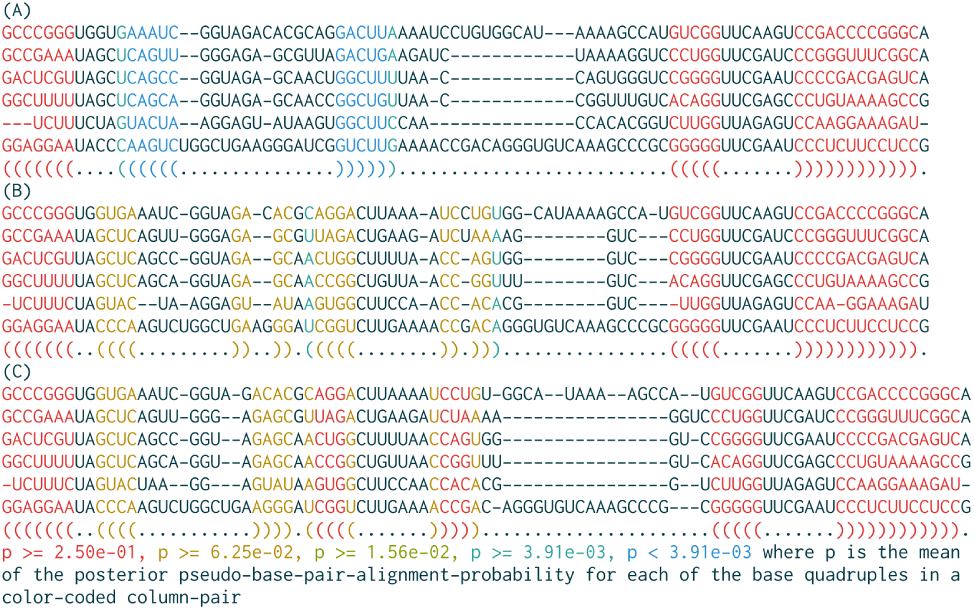
Three examples of a consensus secondary structure color-coded based on pseudo-base-pair-alignment-probabilities 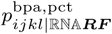 computed by the RNAfamProb program plus the McCaskill program (= the RNAfamProb suite) each with the sequence alignment among six RNA sequences sampled from the tRNA family in the Rfam database. (A) The consensus secondary structure with this alignment computed by the MAFFT G-INS-i program, an maximum-likelihood sequence alignment program (Katoh and Standley, 2013), used for computing this structure by the RNAalifold program, an maximum-likelihood consensus-secondary-structure program (Bernhart et al., 2008). (B) The consensus secondary structure with this alignment computed by the MAFFT X-INS-i program, an maximum-likelihood sequence alignment program considering structural alignment, used for computing this structure by the RNAalifold program. (C) The reference consensus secondary structure derived from the Rfam database with the reference sequence alignment among these six sequences derived from this database. The RNAfamProb program was run with the seven parameter values in Supplementary table 1. A “()” means at least two position pairs in the column pair corresponding to these two parentheses are corresponding.

#### 3.2.2 Visualization of pseudo-base-pairing-probability 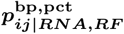 as reliability of two BP positions *i, j*

Six examples of each of 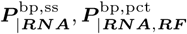 are shown in Figure 6. Two positions *i, j* having 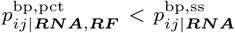 mean the BP by *i, j* is supported less by the consideration of homologous-RNA sequences. On the other hand, two positions *i, j* having 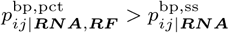 mean the BP by *i, j* is supported more by this consideration. 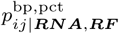 can be used as the reliability of the two BP positions *i, j* as well as 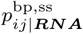 as in Supplementary figure 4.

**Fig. 6.**
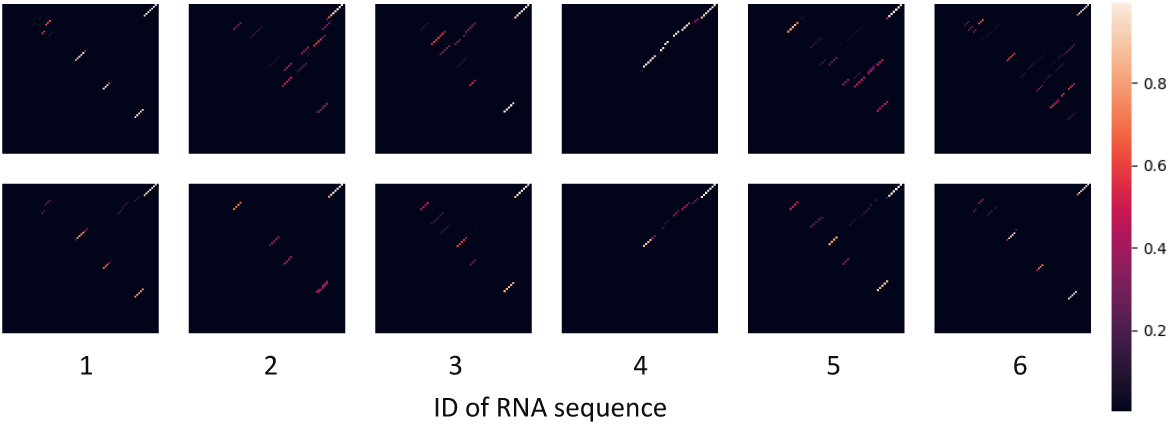
Six examples of each of (upper) a base-pairing probability matrix and (lower) pseudo-base-pairing-probability matrix 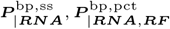. The six examples of a base-pairing probability matrix 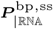 were computed by the McCaskill program. The six examples of a pseudo-base-pairing-probability matrix 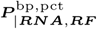 were computed by the RNAfamProb suite with the seven parameter values same as Figure 5. Each of these 12 matrices has the indices 1, 1 at the upper left corner of this matrix. These six sequences are the same as Figure 5.

### 3.3 Comparison of NeoFold algorithm with four maximum-expected-accuracy secondary-structure algorithms

The RNA sequences each having the Rfam database as a source of the ncRNA family to which this sequence belongs and a reference SS of this sequence were collected from the RNA STRAND database, a database of SSs (Andronescu *et al*., 2008), as test set 1. These sequences in this set have no bases except for adenines, guanines, cytosines, and uracils. The SSs in this set have no pseudoknots. Ten RNA sequences and the SS of each of these ten sequences were sampled from a ncRNA family when this family has 11 or more RNA-sequences. The number of the RNA sequences and range of the RNA sequence lengths on each ncRNA family in this set are shown in Supplementary table 2.

The version 0.0.15 of the CentroidFold program provides the CentroidFold and CONTRAfold algorithms. The version 0.0.15 of the CentroidHomFold program provides the CentroidHomFold algorithm. The version 6.0.1 of the TurboFold-smp program provides the TurboFold-MEA algorithm, an MEA SS algorithm with incorporating homologous-RNA sequences via the iterative refinement of BPPs on SS with the help of ML pairwise SAs, with multi-threading. (Tan *et al*., 2017) The principle of each of the CentroidHomFold and TurboFold-MEA algorithms is shown in Supplementary section 7. A distinct difference between these two algorithms and the NeoFold algorithm is whether posterior BPPs for computing MEA SSs are on STA or not.

Each of a positive predictive value (= PPV), sensitivity, and false positive rate (= FPR) is defined as each of

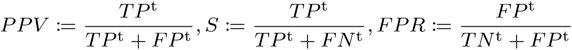

where each of *TP*^t^, *TN*^t^, *FP*^t^, *FN*^t^ is defined as the number of each of TPs, TNs, FPs, and FNs on a test with test set 1. A TP on this test is two positions BP on both two predicted and reference SSs. A TN on this test is a position unpaired on both two predicted and reference SSs. An FP on this test is a position BP on a predicted SS but unpaired on a reference SS. An FN on this test is a position unpaired on a predicted SS but BP on a reference SS. These four numbers are configured in the same way as *a*_***SS*** 𝕊𝕋𝔸_.

The plot of PPVs versus sensitivities by varing the value of a parameter determining the balance between positive and negative (e.g., *γ* on the NeoFold program) among *W* := {2^*w*^|*w* ∈ {−7, …, 10}} for each of the RNAfamProb program plus the McCaskill and NeoFold programs (= the NeoFold suite), and CentroidFold, CentroidHomFold, and TurboFold-smp programs on test set 1 is shown in Figure 7. The plot of FPRs versus sensitivities by varing this value among *W* for each of these suite and three programs on test set 1 is shown in Figure 8. The more a PPV-versus-sensitivity curve faces the upper right corner of the plot space of this curve, the better the prediction accuracy of the estimator corresponding to this curve is. Similarly, the more a FPR-versus-sensitivity (or receiver operating characteristic = ROC) curve faces the upper left corner of the plot space of this curve, the better the prediction accuracy of the estimator corresponding to this curve is. The running time and approximate time-complexity of each of these suite and three programs on test set 1 is shown in Table 1.

**Table 1.** The running time and approximate time-complexity on an RNA family ***RF*** of each of the NeoFold suite and CentroidFold, CentroidHomFold, and TurboFold-smp programs on test set 1. These suite and three programs were run for the value of a parameter determining the balance between positive and negative 2^0^ = 1. These suite and three programs were run on the “Intel Xeon CPU” CPU with 20 CPU logical cores and the clock rate 2.30 [GHz] and 32 [GB] of RAM with the parameter values same as Figure 7. The NeoFold suite and TurboFold-smp program were run with employing multi-threading as much as possible for achieving running time as fast as possible. The CentroidFold and CentroidHomFold programs were run with employing multi-processing as much as possible for achieving running time as fast as possible.

| Program/Suite | Running time [s] | Time complexity |
| --- | --- | --- |
| NeoFold | 31652.036268 | $O(RN^2(R + N))$ |
| CentroidFold (CentroidFold) | 8.569625 | $O(RN^3)$ |
| CentroidFold (CONTRAFold) | 8.698287 | $O(RN^3)$ |
| CentroidHomFold | 99.557175 | $O(R^2N^3)$ |
| TurboFold-smp | 485.652050 | $O(RN^2(R + N))$ |

**Fig. 7.**
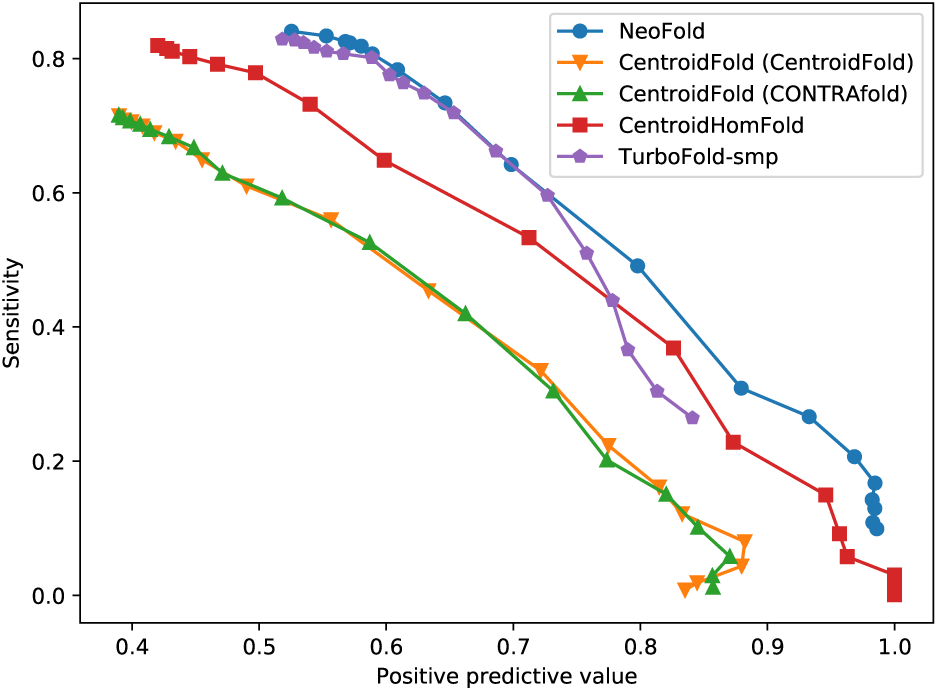
The plot of positive predictive values versus sensitivities by varing the value of a parameter determining the balance between positive and negative among *W* for each of (A) the NeoFold suite and (B) CentroidFold, (C) CentroidHomFold, and (D) TurboFold-smp programs on test set 1. (A) The RNAfamProb program was run with the seven parameter values same as Figure 5. (B, C) The CentroidFold and CentroidHomFold programs were run with the default parameter values of these two programs. The name in a “()” is of the algorithm run by the CentroidFold program. (D) The TurboFold-smp program was run with the number of times of the TurboFold iteration *η* ← 1 and default values of the parameters other than *η*.

**Fig. 8.**
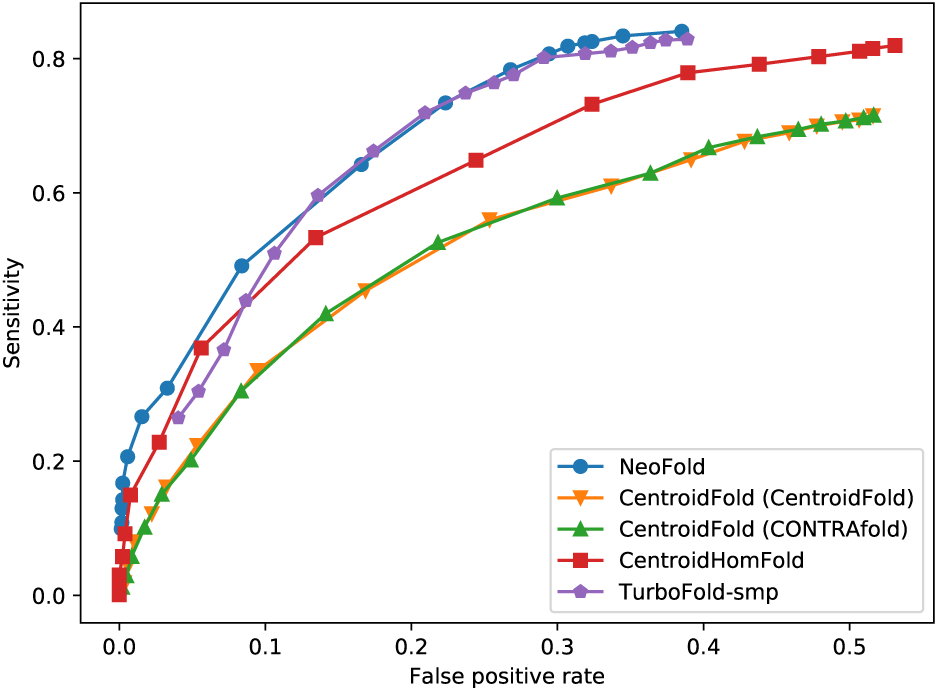
The plot of false positive rates versus sensitivities by varing the value of a parameter determining the balance between positive and negative among *W* for each of (A) the NeoFold suite and (B) CentroidFold, (C) CentroidHomFold, and (D) TurboFold-smp programs on test set 1. These suite and three programs were run with the parameter values same as Figure 7.

The ascending order of the number of PFs computed by each of these suite and three programs is the CentroidFold program → the CentroidHomFold and TurboFold-smp programs → the NeoFold suite. Therefore, the ascending order of the complexities of the probabilities computed by each of these suite and three programs is the same as this order of the numbers of these PFs. Thus, it is expected the NeoFold suite is the best of these suite and three programs in terms of prediction accuracy but worst of these suite and three programs in terms of running time. The prediction accuracy ascending order of these suite and three programs is observed to be the CentroidFold program → the CentroidHomFold prgoram → the TurboFold-smp program → the NeoFold suite, as expected. On the other hand, the running time ascending order of these suite and three programs is observed to be the CentroidFold → program the CentroidHomFold program → the TurboFold-smp program → the NeoFold suite, as expected.

## 4 Conclusions

The RNAfamProb algorithm, an algorithm for estimating pseudo-probabilities given RNA sequences on STA, and NeoFold algorithm, an MEA SS algorithm with these pseudo-probabilities, have been invented. The RNAfamProb and NeoFold programs, an implementation of each of the RNAfamProb and NeoFold algorithms, demonstrated prediction accuracy better than three state-of-the-art MEA-SS-programs while demanding running time far longer than these three programs as expected due to the intrinsic serious problem-complexity of STA compared with independent SS and SA.

The RNAfamProb algorithm can be immediately applied to the CentroidAlign algorithm, CentroidAliFold algorithm, and DAFS algorithm for improving the prediction accuracy of these three algorithms.

Even a novel type of MEA CSS algorithm can be invented with the RNAfamProb algorithm. A CSS between a sequence pair ℝℕ𝔸 is defined as ***CSS***_ℝℕ𝔸_ := (*css*_*ijkl*_) where *css*_*ijkl*_ is defined as 1 if the two position pairs [*i, j*], [*k, l*] are corresponding and 0 otherwise. An example of a CSS ***CSS***_ℝℕ𝔸_ is shown on (A) in Figure 9. The MEA CSS between ℝℕ𝔸, 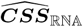, can be computed according to

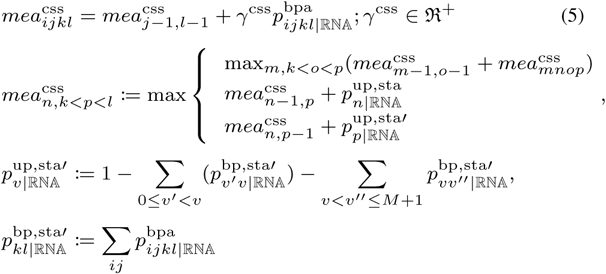

where 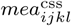 is defined as the EA of 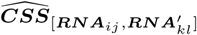. An interpretation of each of the formulation of 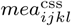 and definition of 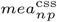 is shown on each of (B) and (C) in Figure 9. 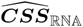 with Equation 5 can be gained without any ***SA***_ℝℕ𝔸_, required by CSS programs so far such as the RNAalifold and CentroidAliFold programs. The quality of CSSs computed from these CSS programs has heavily depended on the quality of SAs supplied with these CSS programs for identifying candidates of two corresponding position-pairs [*i, j*], [*k, l*]. Therefore, the quality of MEA CSSs with Equation 5 will be generally superior to the quality of CSSs computed with these CSS programs.

**Fig. 9.**
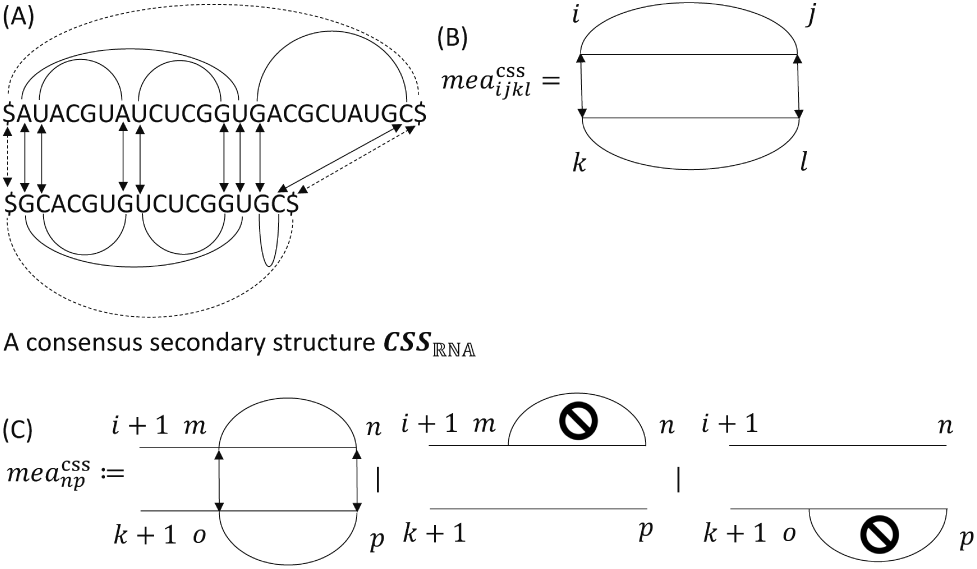
(A) An example of a consensus secondary structureCSS_ℝℕ𝔸_ and interpretation of each of (A) the formulation of maximum expected accuracy 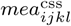 and (B) definition of maximum expected accuracy 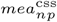.

The RNAfamProb algorithm plus the NeoFold algorithm will be able to estimate more-accurate SSs by incorporating SS probing data such as data from the icSHAPE (= one of experiments probing unpaired bases) (Spitale *et al*., 2015) of homologous-RNA sequences like the SuperFold algorithm, an SS algorithm with incorporating data from the SHAPE-MaP (= one of experiments probing unpaired bases) (Siegfried *et al*., 2014).

## Acknowledgements

We thank the members of the Asai and Frith laboratories for discussing this study with us. Computations were partially performed on the NIG supercomputer at the ROIS National Institute of Genetics.

