## Supplementary Materials for "RNAfamProb Plus NeoFold: Estimations of Posterior Probabilities on RNA Structural Alignment and RNA Secondary Structures with Incorporating Homologous-RNA Sequences"

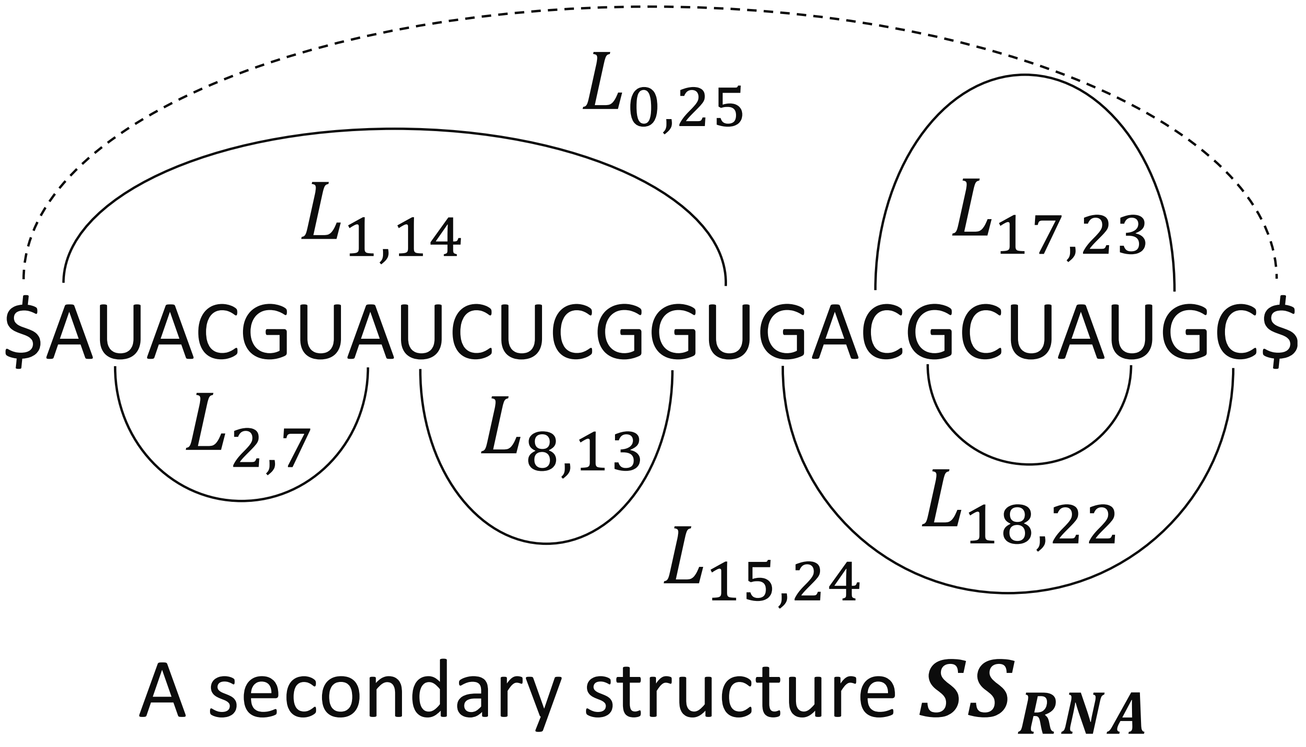


**Fig. 1.** An example of a secondary structure $\boldsymbol{SS_{RNA}}$ with loops $L_{ij}$. $L_{0,25} = \{1, 14, 15, 24\}$. $L_{1,14} = \{2, 7, 8, 13\}$. $L_{15,24} = \{16, 17, 23\}$, $L_{17,23} = \{18, 22\}$. $L_{2,7} = \{3, \ldots, 6\}$, $L_{8,13} = \{9, \ldots, 12\}$, $L_{18,22} = \{19 \ldots, 21\}$. (A) Each of $L_{0,25} = \{1, 14, 15, 24\}$, $L_{1,14} = \{2, 7, 8, 13\}$ is a multi-loop. Each of $L_{15,24} = \{16, 17, 23\}$, $L_{17,23} = \{18, 22\}$ is a 2-loop. Each of $L_{2,7} = \{3, \ldots, 6\}$, $L_{8,13} = \{9, \ldots, 12\}$, $L_{18,22} = \{19 \ldots, 21\}$ is a 1-loop.

#### 1 Recursive formulations of partition functions for inside step of Algorithm 1

A position $u$ is *accessible from two BP positions* $i, j$ if and only if $\exists uij(i < u < j)$ is true and the BP position-pair $[i, j]$ is the closest to $u$ of all BP position-pairs. $L_{ij} := \{u | I^{1}_{uij} = 1\}$ is the *loop* of the two BP positions $i, j$ where $I^{1}_{uij}$ is defined as 1 if the position $u$ is accessible from $i, j$ and 0 otherwise. $L_{ij}$ is *accessible from a loop* $L_{i''j''}; 0 \leq i' < j' \leq N + 1$ if and only if $i, j \in L_{i'j'}$ is true. $L_{ij}$ is a *1-loop* (= a *harpin loop*) if and only if there are no loops accessible from $L_{ij}$ between the positions $i + 1, j - 1$; *2-loop* if and only if there is one loop accessible from $L_{ij}$ between $i + 1, j - 1$; or *multi-loop* if and only if there are more than one loops accessible from $L_{ij}$ between $i + 1, j - 1$. An example of $\boldsymbol{SS_{RNA}}$ with $L_{ij}$ is shown in Supplementary figure 1.

$fe_{\boldsymbol{SS}}$ can be formulated as

$$fe_{\boldsymbol{SS}} = \sum_{ij, \lambda \in \{1\mathrm{l}, 2\mathrm{l}, \mathrm{ml}\} | \mathrm{I}^{\mathrm{bp}}_{\mathrm{SSij}} \mathrm{I}^{\lambda}_{\mathrm{ij}} = 1} fe^{\lambda}_{ij}$$

where each of $I^{1\mathrm{l}}_{\boldsymbol{SS}ij}, I^{2\mathrm{l}}_{\boldsymbol{SS}ij}, I^{\mathrm{ml}}_{\boldsymbol{SS}ij}$ is defined as 1 if $L_{ij}$ is each of a 1-loop, 2-loop, and multi-loop in $\boldsymbol{SS}$ and 0 otherwise and each of $fe^{1\mathrm{l}}_{ij}, fe^{2\mathrm{l}}_{ij}, fe^{\mathrm{ml}}_{ij}$ the FE of $L_{ij}$ being each of these three loops. $fe^{2\mathrm{l}}_{ij}$ can be formulated as

$$fe^{2\mathrm{l}}_{ij} = \sum_{mn | I^{\mathrm{bp}}_{\boldsymbol{SS}mn} I^{1}_{mij} I^{1}_{nij} = 1} fe^{2\mathrm{l}}_{ijmn}$$

where $fe^{2\mathrm{l}}_{ijmn}$ is defined as the FE of $L_{ij}$ containing $L_{mn}$ accessible from $L_{ij}$. $fe^{\mathrm{ml}}_{ij}$ can be formulated as

$$fe^{\mathrm{ml}}_{ij} = fe^{\mathrm{ml,cbp}} + \sum_{mn | I^{\mathrm{bp}}_{\boldsymbol{SS}mn} I^{1}_{mij} I^{1}_{nij} = 1} fe^{\mathrm{ml,abp}}; fe^{\mathrm{ml,cbp}}, fe^{\mathrm{ml,abp}} \in \Re$$

where $\Re$ is defined as the real-number space. Each of $fe^{\mathrm{ml,cbp}}, fe^{\mathrm{ml,abp}}$ is each of multi-loop offset and per-accessible-BP FE. $fe^{1\mathrm{l}}_{ij}, fe^{2\mathrm{l}}_{ijmn}, fe^{\mathrm{ml,cbp}}, fe^{\mathrm{ml,abp}}$ can be computed by the Turner model (Turner and Mathews, 2010).

$Z^{\mathrm{sta,bpa,1l}}_{ijkl}$ is defined as

$$Z^{\mathrm{sta,bpa,1l}}_{ijkl} = (or^{\mathrm{bpa}}_{ijkl})^{\beta^{\mathrm{sta}} s} \exp(-\beta^{\mathrm{sta}}(fe^{1\mathrm{l}}_{ij} + fe^{1\mathrm{l}}_{kl})) Z^{\mathrm{sa,f}}_{j-1,l-1};$$

$$or^{\mathrm{bpa}}_{ijkl} := \begin{cases} 1 & (i = k = 0 \land j = N + 1 \land l = M + 1) \\ \frac{p^{\mathrm{bpa}}_{ijkl}}{p^{\mathrm{bpa,r}}_{ijkl}} & (otherwise) \end{cases}$$

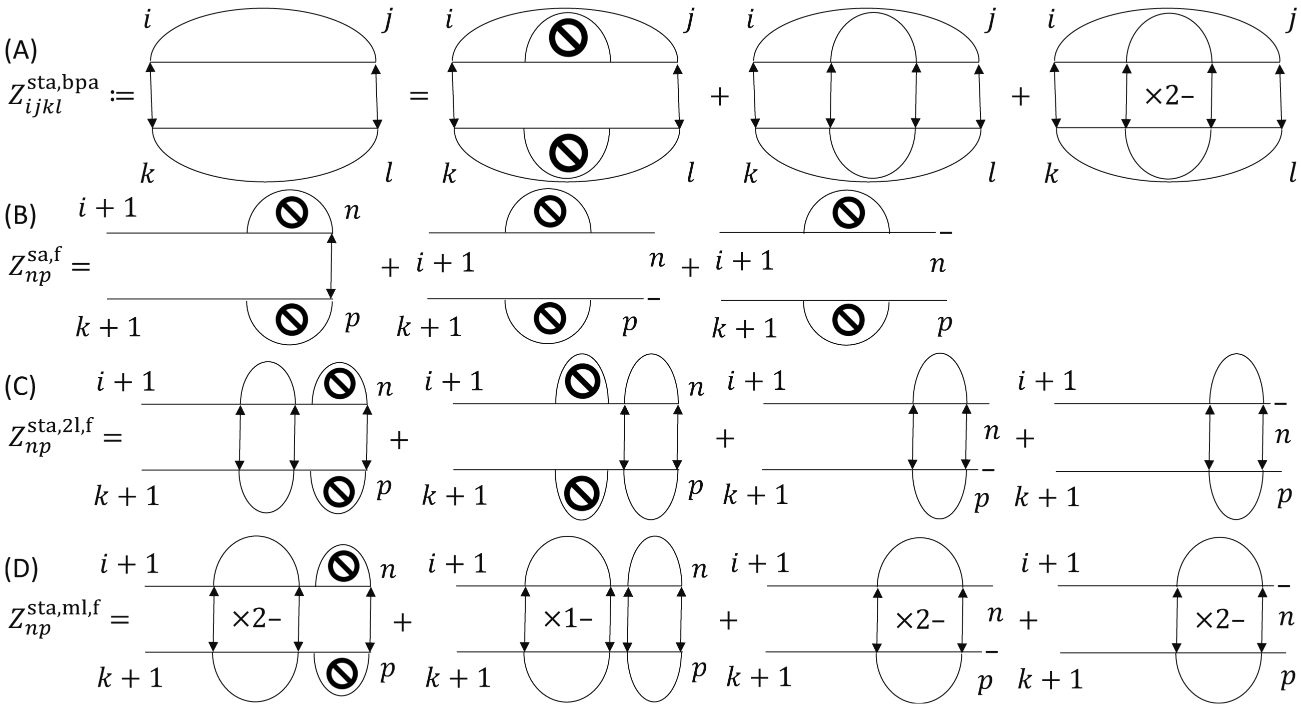


**Fig. 2.** An interpretation of the formulation of each of four partition functions (A)$Z^{\text{sta,bpa}}_{ijkl}$, (B)$Z^{\text{sa,f}}_{np}$, (C)$Z^{\text{sta,2l,f}}_{np}$, (D)$Z^{\text{sta,ml,f}}_{np}$.

where $Z^{\text{sa,f}}_{np}$ is defined as the PF on $\mathcal{STA}_{\mathbb{RNA}_{i+1,n,k+1,p}}$ in the case in which each of $L_{ij}, L_{kl}$ is a 1-loop. $Z^{\text{sta,bpa,2l}}_{ijkl}$ is defined as

$$Z^{\text{sta,bpa,2l}}_{ijkl} = (or^{\text{bpa}}_{ijkl})^{\beta^{\text{sta}} s} Z^{\text{sta,2l,f}}_{j-1,l-1}$$

where $Z^{\text{sta,2l,f}}_{np}$ is defined as the PF on $\mathcal{STA}_{\mathbb{RNA}_{i+1,n,k+1,p}}$ in the case in which each of $L_{ij}, L_{kl}$ is a 2-loop multiplied by $\sum_{mn}(exp(-\beta^{\text{sta}} fe^{\text{2l}}_{ijmn})) + \sum_{op} \exp(-\beta^{\text{sta}} fe^{\text{2l}}_{klop})$. $Z^{\text{sta,bpa,ml}}_{ijkl}$ is defined as

$$Z^{\text{sta,bpa,ml}}_{ijkl} = (or^{\text{bpa}}_{ijkl})^{\beta^{\text{sta}} s} \exp(-2\beta^{\text{sta}} fe^{\text{ml,cbp}}) Z^{\text{sta,ml,f}}_{j-1,l-1}$$

where $Z^{\text{sta,ml,f}}_{np}$ is defined as the PF on $\mathcal{STA}_{\mathbb{RNA}_{i+1,n,k+1,p}}$ in the case in which each of $L_{ij}, L_{kl}$ is a multi-loop. $Z^{\text{sta,bpa}}_{ijkl}$ is defined as the PF on $\mathfrak{STA}_{[\boldsymbol{RNA}_{ij}, \boldsymbol{RNA}'_{kl}]}$ in the case in which the two position pairs $[i, j]$, $[k, l]$ are corresponding. $Z^{\text{sta}}$ is gained from the fact $Z^{\text{sta}}$, $Z^{\text{sta,bpa}}_{0,N+1,0,M+1}$ are equal for any $\mathbb{RNA}$. $Z^{\text{sta,bpa}}_{ijkl}$ can be formulated as

$$Z^{\text{sta,bpa}}_{ijkl} = Z^{\text{sta,bpa,1l}}_{ijkl} + Z^{\text{sta,bpa,2l}}_{ijkl} + Z^{\text{sta,bpa,ml}}_{ijkl}.$$

An interpretation of the formulation of $Z^{\text{sta,bpa}}_{ijkl}$ is shown on (A) in Supplementary figure 2.

#### 1.1 Formulation of partition function $Z^{\text{sa,f}}_{np}$

$Z^{\text{sa,f}}_{np}$ can be formulated as

$$Z^{\text{sa,f}}_{np} = Z^{\text{sa,f,ba}}_{np} + Z^{\text{sa,f,g}}_{np} + Z^{\text{sa,f,g}\prime}_{np}$$

where $Z^{\text{sa,f,ba}}_{np}$ is defined as $Z^{\text{sa,f}}_{np}$ in the case in which the two positions $n, p$ are aligned; $Z^{\text{sa,f,og}}_{np}$ $Z^{\text{sa,f}}_{np}$ in the case in which $n$ is aligned with a gap behind $p$; and $Z^{\text{sa,f,og}\prime}_{np}$ $Z^{\text{sa,f}}_{np}$ in the case in which $p$ is aligned with a gap behind $n$. $Z^{\text{sa,f,ba}}_{np}$ can be formulated as

$$Z^{\text{sa,f,ba}}_{np} = Z^{\text{sa,f}}_{n-1,p-1}(or^{\text{ba}}_{np})^{\beta^{\text{sta}} s}; or^{\text{ba}}_{uv} := \frac{p^{\text{ba,sta}}_{uv}}{p^{\text{ba,sta,r}}_{uv}}.$$

$Z^{\text{sa,f,g}}_{np}$ can be formulated as

$$Z^{\text{sa,f,g}}_{np} = (Z^{\text{sa,f,ba}}_{n-1,p} + Z^{\text{sa,f,g}\prime}_{n-1,p}) \exp(\beta^{\text{sta}} sp^{\text{og}}) + Z^{\text{sa,f,g}}_{n-1,p} \exp(\beta^{\text{sta}} sp^{\text{eg}}).$$

$Z^{\text{sa,f,g}\prime}_{np}$ can be formulated as

$$Z^{\text{sa,f,g}\prime}_{np} = (Z^{\text{sa,f,ba}}_{n,p-1} + Z^{\text{sa,f,og}}_{n,p-1}) \exp(\beta^{\text{sta}} sp^{\text{og}}) + Z^{\text{sa,f,g}\prime}_{n,p-1} \exp(\beta^{\text{sta}} sp^{\text{eg}}).$$

Each of $Z^{\text{sa,f}}_{ik}, Z^{\text{sa,f,ba}}_{ik}, Z^{\text{sa,f,g}}_{ik}, Z^{\text{sa,f,g}\prime}_{ik}$ is formulated (or initialized) as each of

$$Z^{\text{sa,f}}_{ik} = Z^{\text{sa,f,ba}}_{ik} = 1, Z^{\text{sa,f,g}}_{ik} = Z^{\text{sa,f,g}\prime}_{ik} = 0.$$

##### 1.2 Formulation of partition function $Z_{np}^{\mathrm{sta,2l,f}}$

$Z_{np}^{\mathrm{sta,2l,f}}$ can be formulated as

$$Z_{np}^{\mathrm{sta,2l,f}} = Z_{np}^{\mathrm{sta,2l,f,ba}} + Z_{np}^{\mathrm{sta,2l,f,g}} + Z_{np}^{\mathrm{sta,2l,f,g\prime}}$$

where $Z_{np}^{\mathrm{sta,2l,f,ba}}$ is defined as $Z_{np}^{\mathrm{sta,2l,f}}$ in the case in which the two positions $n, p$ are aligned; $Z_{np}^{\mathrm{sta,2l,f,g}}$ $Z_{np}^{\mathrm{sta,2l,f}}$ in the case in which $n$ is aligned with a gap behind $p$; and $Z_{np}^{\mathrm{sta,2l,f,g\prime}}$ $Z_{np}^{\mathrm{sta,2l,f}}$ in the case in which $p$ is aligned with a gap behind $n$. $Z_{np}^{\mathrm{sta,2l,f,ba}}$ can be formulated as

$$Z_{np}^{\mathrm{sta,2l,f,ba}} = \sum_{mo} [Z_{m-1,o-1}^{\mathrm{sa,f}} \exp(-\beta^{\mathrm{sta}}(fe_{ijmn}^{\mathrm{2l}} + fe_{klop}^{\mathrm{2l}})) Z_{mnop}^{\mathrm{sta,bpa}}] + Z_{n-1,p-1}^{\mathrm{sta,2l,f}} (or_{np}^{\mathrm{ba}})^{\beta^{\mathrm{sta}} s}.$$

$Z_{np}^{\mathrm{sta,2l,f,g}}$ can be formulated as

$$Z_{np}^{\mathrm{sta,2l,f,og}} = (Z_{n-1,p}^{\mathrm{sta,2l,f,ba}} + Z_{n-1,p}^{\mathrm{sta,2l,f,g\prime}}) \exp(\beta^{\mathrm{sta}} sp^{\mathrm{og}}) + Z_{n-1,p}^{\mathrm{sta,2l,f,g}} \exp(\beta^{\mathrm{sta}} sp^{\mathrm{eg}}).$$

$Z_{np}^{\mathrm{sta,2l,f,g\prime}}$ can be formulated as

$$Z_{np}^{\mathrm{sta,2l,f,og\prime}} = (Z_{n,p-1}^{\mathrm{sta,2l,f,ba}} + Z_{n,p-1}^{\mathrm{sta,2l,f,g}}) \exp(\beta^{\mathrm{sta}} sp^{\mathrm{og}}) + Z_{n,p-1}^{\mathrm{sta,2l,f,g\prime}} \exp(\beta^{\mathrm{sta}} sp^{\mathrm{eg}}).$$

Each of $Z_{ik}^{\mathrm{sta,2l,f}}, Z_{ik}^{\mathrm{sta,2l,f,ba}}, Z_{ik}^{\mathrm{sta,2l,f,g}}, Z_{ik}^{\mathrm{sta,2l,f,g\prime}}$ is formulated (or initialized) as each of

$$Z_{ik}^{\mathrm{sta,2l,f}} = Z_{ik}^{\mathrm{sta,2l,f,ba}} = Z_{ik}^{\mathrm{sta,2l,f,g}} = Z_{ik}^{\mathrm{sta,2l,f,g\prime}} = 0.$$

##### 1.3 Formulation of partition function $Z_{np}^{\mathrm{sta,ml,f}}$

$Z_{np}^{\mathrm{sta,ml,f}}$ can be formulated as

$$Z_{np}^{\mathrm{sta,ml,f}} = Z_{np}^{\mathrm{sta,ml,f,ba}} + Z_{np}^{\mathrm{sta,ml,f,g}} + Z_{np}^{\mathrm{sta,ml,f,g\prime}}$$

where $Z_{np}^{\mathrm{sta,ml,f,ba}}$ is defined as $Z_{np}^{\mathrm{sta,ml,f}}$ in the case in which the two positions $n, p$ are aligned; $Z_{np}^{\mathrm{sta,ml,f,g}}$ $Z_{np}^{\mathrm{sta,ml,f}}$ in the case in which $n$ is aligned with a gap behind $p$; and $Z_{np}^{\mathrm{sta,ml,f,g\prime}}$ $Z_{np}^{\mathrm{sta,ml,f}}$ in the case in which $p$ is aligned with a gap behind $n$. $Z_{np}^{\mathrm{sta,ml,f,ba}}$ can be formulated as

$$Z_{np}^{\mathrm{sta,ml,f,ba}} = \sum_{mo} [(Z_{m-1,o-1}^{\mathrm{sta,abp,f}} + Z_{m-1,o-1}^{\mathrm{sta,ml,f}}) \exp(-2\beta^{\mathrm{sta}} fe^{\mathrm{ml,abp}}) Z_{mnop}^{\mathrm{sta,bpa}}] + Z_{n-1,p-1}^{\mathrm{sta,ml,f}} (or_{np}^{\mathrm{ba}})^{\beta^{\mathrm{sta}} s}$$

where $Z_{np}^{\mathrm{sta,abp,f}}$ is defined as $Z_{np}^{\mathrm{sta,ml,f}}$ in the case in which each of $L_{ij}, L_{kl}$ contains two BP positions between each of the two position pairs $[i+1, n]$, $[k+1, p]$. $Z_{np}^{\mathrm{sta,ml,f,og}}$ can be formulated as

$$Z_{np}^{\mathrm{sta,ml,f,g}} = (Z_{n-1,p}^{\mathrm{sta,ml,f,ba}} + Z_{n-1,p}^{\mathrm{sta,ml,f,g\prime}}) \exp(\beta^{\mathrm{sta}} sp^{\mathrm{og}}) + Z_{n-1,p}^{\mathrm{sta,ml,f,g}} \exp(\beta^{\mathrm{sta}} sp^{\mathrm{eg}}).$$

$Z_{np}^{\mathrm{sta,ml,f,g\prime}}$ can be formulated as

$$Z_{np}^{\mathrm{sta,ml,f,g\prime}} = (Z_{n,p-1}^{\mathrm{sta,ml,f,ba}} + Z_{n,p-1}^{\mathrm{sta,ml,f,og}}) \exp(\beta^{\mathrm{sta}} sp^{\mathrm{og}}) + Z_{n,p-1}^{\mathrm{sta,ml,f,g\prime}} \exp(\beta^{\mathrm{sta}} sp^{\mathrm{eg}}).$$

Each of $Z_{ik}^{\mathrm{sta,ml,f}}, Z_{ik}^{\mathrm{sta,ml,f,ba}}, Z_{ik}^{\mathrm{sta,ml,f,g}}, Z_{ik}^{\mathrm{sta,ml,f,g\prime}}$ is formulated (or initialized) as each of

$$Z_{ik}^{\mathrm{sta,ml,f}} = Z_{ik}^{\mathrm{sta,ml,f,ba}} = Z_{ik}^{\mathrm{sta,ml,f,g}} = Z_{ik}^{\mathrm{sta,ml,f,g\prime}} = 0.$$

$Z_{np}^{\mathrm{sta,abp,f}}$ can be formulated as

$$Z_{np}^{\mathrm{sta,abp,f}} = Z_{np}^{\mathrm{sta,abp,f,ba}} + Z_{np}^{\mathrm{sta,abp,f,g}} + Z_{np}^{\mathrm{sta,abp,f,g\prime}}$$

where $Z_{np}^{\mathrm{sta,abp,f,ba}}$ is defined as $Z_{np}^{\mathrm{sta,abp,f}}$ in the case in which the two positions $n, p$ are aligned; $Z_{np}^{\mathrm{sta,abp,f,g}}$ $Z_{np}^{\mathrm{sta,abp,f}}$ in the case in which $n$ is aligned with a gap behind $p$; and $Z_{np}^{\mathrm{sta,abp,f,g\prime}}$ $Z_{np}^{\mathrm{sta,abp,f}}$ in the case in which $p$ is aligned with a gap behind $n$. $Z_{np}^{\mathrm{sta,abp,f,ba}}$ can be formulated as

$$Z_{np}^{\mathrm{sta,abp,f,ba}} = \sum_{mo} (Z_{m-1,o-1}^{\mathrm{sa,f}} \exp(-2\beta^{\mathrm{sta}} fe^{\mathrm{ml,abp}}) Z_{mnop}^{\mathrm{sta,bpa}}) + Z_{n-1,p-1}^{\mathrm{sta,abp,f}} (or_{np}^{\mathrm{ba}})^{\beta^{\mathrm{sta}} s}.$$

$Z_{np}^{\mathrm{sta,abp,f,g}}$ can be formulated as

$$Z_{np}^{\mathrm{sta,abp,f,g}} = (Z_{n-1,p}^{\mathrm{sta,abp,f,ba}} + Z_{n-1,p}^{\mathrm{sta,abp,f,g\prime}}) \exp(\beta^{\mathrm{sta}} sp^{\mathrm{og}}) + Z_{n-1,p}^{\mathrm{sta,abp,f,g}} \exp(\beta^{\mathrm{sta}} sp^{\mathrm{eg}}).$$

$Z_{np}^{\mathrm{sta,abp,f,g\prime}}$ can be formulated as

$$Z_{np}^{\mathrm{sta,abp,f,g\prime}} = (Z_{n,p-1}^{\mathrm{sta,abp,f,ba}} + Z_{n,p-1}^{\mathrm{sta,abp,f,g}}) \exp(\beta^{\mathrm{sta}} sp^{\mathrm{og}}) + Z_{n,p-1}^{\mathrm{sta,abp,f,g\prime}} \exp(\beta^{\mathrm{sta}} sp^{\mathrm{eg}}).$$

Each of $Z_{ik}^{\mathrm{sta,abp,f}}, Z_{ik}^{\mathrm{sta,abp,f,ba}}, Z_{ik}^{\mathrm{sta,abp,f,g}}, Z_{ik}^{\mathrm{sta,abp,f,g\prime}}$ is formulated (or initialized) as each of

$$Z_{ik}^{\mathrm{sta,abp,f}} = Z_{ik}^{\mathrm{sta,abp,f,ba}} = Z_{ik}^{\mathrm{sta,abp,f,g}} = Z_{ik}^{\mathrm{sta,abp,f,g\prime}} = 0.$$

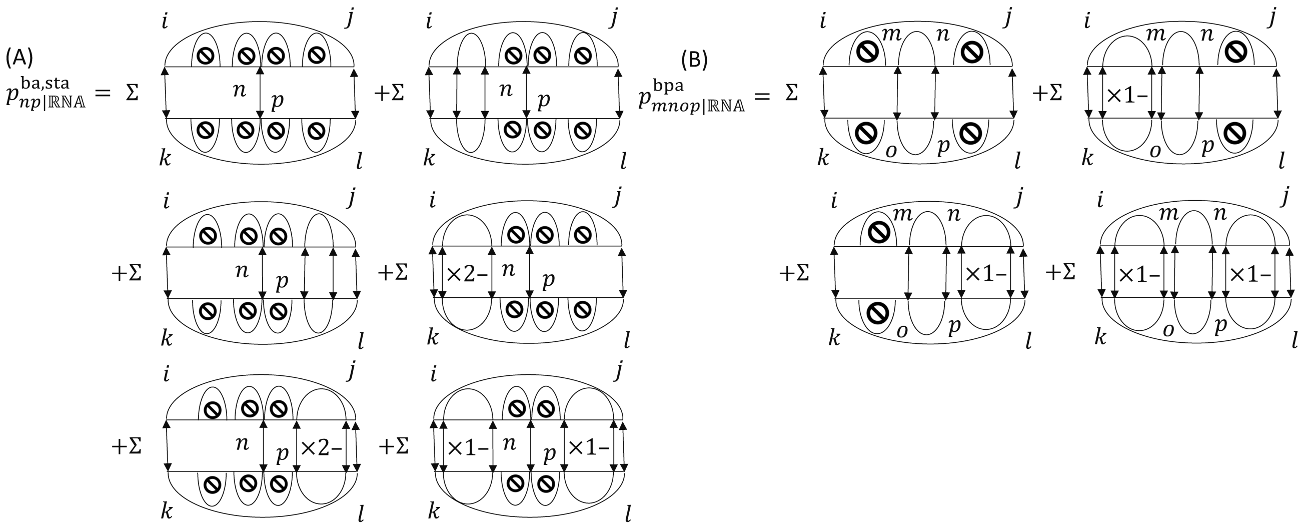


**Fig. 3.** An interpretation of the formulation of each of a base alignment probability and base pair alignment probability (A)$p^{\mathrm{ba,sta}}_{np|\mathbb{RNA}}$, (B)$p^{\mathrm{bpa}}_{mnop|\mathbb{RNA}}$.

##### 1.4 Analysis of time and space complexities of computing partition functions $Z^{\mathrm{sta,bpa}}_{ijkl}$

Lemma 1.1. $Z^{\mathrm{sta,bpa}}_{ijkl}$ *can be computed on a DP with the time and space complexities* $O(N^4M^4), O(N^2M^2)$.

Proof. $Z^{\mathrm{sta,bpa}}_{ijkl}, Z^{\mathrm{sta,bpa},\lambda}_{ijkl}$ can be computed on a DP with the time and space complexities $O(1), O(1)$ since each of $Z^{\mathrm{sta,bpa}}_{ijkl}, Z^{\mathrm{sta,bpa},\lambda}_{ijkl}$ involves only the four positions $i, j, k, l$ in the formulation of each of $Z^{\mathrm{sta,bpa}}_{ijkl}, Z^{\mathrm{sta,bpa},\lambda}_{ijkl}$. $Z^{\cdot}_{np}$ can be computed on a DP with at the most time and space complexities $O(NM), O(NM)$ since $Z^{\cdot}_{np}$ involves at most positions $m, o$ and the two positions $n, p$ in the formulation of $Z^{\cdot}_{np}$. $Z^{\cdot}_{np}$ can be discarded after $Z^{\mathrm{sta,bpa}}_{ijkl}, Z^{\mathrm{sta,bpa},\lambda}_{ijkl}$ are computed. Therefore, $Z^{\mathrm{sta,bpa}}_{ijkl}$ can be computed on a DP with the time and space complexities $O(N^4M^4), O(N^2M^2)$.

DPs like this DP are called inside algorithms since recursions for these DPs are proceeded inside.

#### 2 Recursive formulation of each of base alignment probability and base pair alignment probability $p^{\mathrm{ba,sta}}_{uv|\mathbb{RNA}}, p^{\mathrm{bpa}}_{ijkl|\mathbb{RNA}}$ for outside step of Algorithm 1

##### 2.1 Formulation of base alignment probability $p^{\mathrm{ba,sta}}_{np|\mathbb{RNA}}$

$Z^{\mathrm{sa,b}}_{mo}$ is defined as the PF on $\mathcal{STA}_{\mathbb{RNA}_{m,j-1,o,l-1}}$ in the case in which each of $L_{ij}, L_{kl}$ is a 1-loop. $Z^{\mathrm{sta,2l,b}}_{mo}$ is defined as the PF on $\mathcal{STA}_{\mathbb{RNA}_{m,j-1,o,l-1}}$ in the case in which each of $L_{ij}, L_{kl}$ is a 2-loop multiplied by $\sum_{mn}(exp(-\beta^{\mathrm{sta}} fe^{\mathrm{2l}}_{ijmn})) + \sum_{op}\exp(-\beta^{\mathrm{sta}} fe^{\mathrm{2l}}_{klop})$. $Z^{\mathrm{sta,ml,b}}_{mo}$ is defined as the PF on $\mathcal{STA}_{\mathbb{RNA}_{m,j-1,o,l-1}}$ in the case in which each of $L_{ij}, L_{kl}$ is a multi-loop. $Z^{\mathrm{sta,abp,b}}_{mo}$ is defined as $Z^{\mathrm{sta,ml,b}}_{mo}$ in the case in which each of $L_{ij}, L_{kl}$ contains two BP positions between each of the two position pairs $[m, j-1], [o, l-1]$. A base alignment probability $p^{\mathrm{ba,sta}}_{np|\mathbb{RNA}}$ can be formulated as

$$p^{\mathrm{ba,sta}}_{np|\mathbb{RNA}} = (or^{\mathrm{ba}}_{np})^{\beta^{\mathrm{sta}}s} \sum_{ijkl} \Big\{ \frac{p^{\mathrm{bpa}}_{ijkl|\mathbb{RNA}} (or^{\mathrm{bpa}}_{ijkl})^{\beta^{\mathrm{sta}}s}}{Z^{\mathrm{sta,bpa}}_{ijkl}} [\exp(-\beta^{\mathrm{sta}}(fe^{\mathrm{1l}}_{ij} + fe^{\mathrm{1l}}_{kl})) Z^{\mathrm{sa,f}}_{n-1,p-1} Z^{\mathrm{sa,b}}_{n+1,p+1} + (Z^{\mathrm{sta,2l,f}}_{n-1,p-1} Z^{\mathrm{sa,b}}_{n+1,p+1} + Z^{\mathrm{sa,f}}_{n-1,p-1} Z^{\mathrm{sta,2l,b}}_{n+1,p+1}) + \exp(-2\beta^{\mathrm{sta}} fe^{\mathrm{ml,cbp}})(Z^{\mathrm{sta,ml,f}}_{n-1,p-1} Z^{\mathrm{sa,b}}_{n+1,p+1} + Z^{\mathrm{sa,f}}_{n-1,p-1} Z^{\mathrm{sta,ml,b}}_{n+1,p+1} + Z^{\mathrm{sta,ml,f}}_{n-1,p-1} Z^{\mathrm{sta,abp,b}}_{n+1,p+1} + Z^{\mathrm{sta,abp,f}}_{n-1,p-1} Z^{\mathrm{sta,ml,b}}_{n+1,p+1} + Z^{\mathrm{sta,ml,f}}_{n-1,p-1} Z^{\mathrm{sta,ml,b}}_{n+1,p+1} + Z^{\mathrm{sta,abp,f}}_{n-1,p-1} Z^{\mathrm{sta,abp,b}}_{n+1,p+1})] \Big\}.$$

An interpretation of the formulation of $p^{\mathrm{ba,sta}}_{np|\mathbb{RNA}}$ is shown on (A) in Supplementary figure 3.

###### 2.1.1 Formulation of $Z^{\mathrm{sa,b}}_{mo}$

$Z^{\mathrm{sa,b}}_{mo}$ can be formulated as

$$Z^{\mathrm{sa,b}}_{mo} = Z^{\mathrm{sa,b,ba}}_{mo} + Z^{\mathrm{sa,b,g}}_{mo} + Z^{\mathrm{sa,b,g\prime}}_{mo}$$

where $Z^{\mathrm{sa,b,ba}}_{mo}$ is defined as $Z^{\mathrm{sa,b}}_{mo}$ in the case in which the two positions $m, o$ are aligned; $Z^{\mathrm{sa,b,g}}_{mo}$ $Z^{\mathrm{sa,b}}_{mo}$ in the case in which $m$ is aligned with a gap in front of $o$; and $Z^{\mathrm{sa,b,g\prime}}_{mo}$ $Z^{\mathrm{sa,b}}_{mo}$ in the case in which $o$ is aligned with a gap in front of $m$. $Z^{\mathrm{sa,b,ba}}_{mo}$ can be formulated as

$$Z^{\mathrm{sa,b,ba}}_{mo} = Z^{\mathrm{sa,b}}_{m+1,o+1}(or^{\mathrm{ba}}_{np})^{\beta^{\mathrm{sta}}s}.$$

$Z^{\mathrm{sa,b,g}}_{mo}$ can be formulated as

$$Z^{\mathrm{sa,b,g}}_{mo} = (Z^{\mathrm{sa,b,ba}}_{m+1,o} + Z^{\mathrm{sa,b,g\prime}}_{m+1,o}) \exp(\beta^{\mathrm{sta}} sp^{\mathrm{og}}) + Z^{\mathrm{sa,b,g}}_{m+1,o} \exp(\beta^{\mathrm{sta}} sp^{\mathrm{eg}}).$$

$Z_{mo}^{\text{sa,b,g}\prime}$ can be formulated as

$$Z_{mo}^{\text{sa,b,g}\prime} = (Z_{m,o+1}^{\text{sa,b,ba}} + Z_{m,o+1}^{\text{sa,b,g}}) \exp(\beta^{\text{sta}} sp^{\text{og}}) + Z_{m,o+1}^{\text{sa,b,g}\prime} \exp(\beta^{\text{sta}} sp^{\text{eg}}).$$

Each of $Z_{jl}^{\text{sa,b}}, Z_{jl}^{\text{sa,b,ba}}, Z_{jl}^{\text{sa,b,g}}, Z_{jl}^{\text{sa,b,g}\prime}$ is formulated (or initialized) as each of

$$Z_{jl}^{\text{sa,b}} = Z_{jl}^{\text{sa,b,ba}} = 1, Z_{jl}^{\text{sa,b,g}} = Z_{jl}^{\text{sa,b,g}\prime} = 0.$$

##### 2.1.2 Formulation of $Z_{mo}^{\text{sta,2l,b}}$

$Z_{mo}^{\text{sta,2l,b}}$ can be formulated as

$$Z_{mo}^{\text{sta,2l,b}} = Z_{mo}^{\text{sta,2l,b,ba}} + Z_{mo}^{\text{sta,2l,b,g}} + Z_{mo}^{\text{sta,2l,b,g}\prime}$$

where $Z_{mo}^{\text{sta,2l,b,ba}}$ is defined as the PF $Z_{mo}^{\text{sta,2l,b}}$ in the case in which the two positions $m, o$ are aligned; $Z_{mo}^{\text{sta,2l,b,g}}$ $Z_{mo}^{\text{sta,2l,b}}$ in the case in which $m$ is aligned with a gap in front of $o$; and $Z_{mo}^{\text{sta,2l,b,g}\prime}$ $Z_{mo}^{\text{sta,2l,b}}$ in the case in which $o$ is aligned with a gap in front of $m$. $Z_{mo}^{\text{sta,2l,b,ba}}$ can be formulated as

$$Z_{mo}^{\text{sta,2l,b,ba}} = \sum_{np} [Z_{n+1,p+1}^{\text{sa,b}} \exp(-\beta^{\text{sta}}(fe_{ijmn}^{\text{2l}} + fe_{klop}^{\text{2l}})) Z_{mnop}^{\text{sta,bpa}}] + Z_{m+1,o+1}^{\text{sta,2l,b}} (or_{mo}^{\text{ba}})^{\beta^{\text{sta}} s}.$$

$Z_{mo}^{\text{sta,2l,b,g}}$ can be formulated as

$$Z_{mo}^{\text{sta,2l,b,g}} = (Z_{m+1,o}^{\text{sta,2l,b,ba}} + Z_{m+1,o}^{\text{sta,2l,b,g}\prime}) \exp(\beta^{\text{sta}} sp^{\text{og}}) + Z_{m+1,o}^{\text{sta,2l,b,g}} \exp(\beta^{\text{sta}} sp^{\text{eg}}).$$

$Z_{mo}^{\text{sta,2l,b,g}\prime}$ can be formulated as

$$Z_{mo}^{\text{sta,2l,b,og}\prime} = (Z_{m,o+1}^{\text{sta,2l,b,ba}} + Z_{m,o+1}^{\text{sta,2l,b,g}}) \exp(\beta^{\text{sta}} sp^{\text{og}}) + Z_{m,o+1}^{\text{sta,2l,b,g}\prime} \exp(\beta^{\text{sta}} sp^{\text{eg}}).$$

Each of $Z_{jl}^{\text{sta,2l,b}}, Z_{jl}^{\text{sta,2l,b,ba}}, Z_{jl}^{\text{sta,2l,b,g}}, Z_{jl}^{\text{sta,2l,b,g}\prime}$ is formulated as each of

$$Z_{jl}^{\text{sta,2l,b}} = Z_{jl}^{\text{sta,2l,b,ba}} = Z_{jl}^{\text{sta,2l,b,g}} = Z_{jl}^{\text{sta,2l,b,g}\prime} = 0.$$

##### 2.1.3 Formulation of $Z_{mo}^{\text{sta,ml,b}}$

$Z_{mo}^{\text{sta,ml,b}}$ can be formulated as

$$Z_{mo}^{\text{sta,ml,b}} = Z_{mo}^{\text{sta,ml,b,ba}} + Z_{mo}^{\text{sta,ml,b,g}} + Z_{mo}^{\text{sta,ml,b,g}\prime}$$

where $Z_{mo}^{\text{sta,ml,b,ba}}$ is defined as the PF $Z_{mo}^{\text{sta,ml,b}}$ in the case in which the two positions $m, o$ are aligned; $Z_{mo}^{\text{sta,ml,b,g}}$ $Z_{mo}^{\text{sta,ml,b}}$ in the case in which $m$ is aligned with a gap in front of $o$; and $Z_{mo}^{\text{sta,ml,b,g}\prime}$ $Z_{mo}^{\text{sta,ml,b}}$ in the case in which $o$ is aligned with a gap in front of $m$. $Z_{mo}^{\text{sta,ml,b,ba}}$ can be formulated as

$$Z_{mo}^{\text{sta,ml,b,ba}} = \sum_{np} [(Z_{n+1,p+1}^{\text{sta,abp,b}} + Z_{n+1,p+1}^{\text{sta,ml,b}}) \exp(-2\beta^{\text{sta}} fe^{\text{ml,abp}}) Z_{mnop}^{\text{sta,bpa}}] + Z_{m+1,o+1}^{\text{sta,ml,b}} (or_{mo}^{\text{ba}})^{\beta^{\text{sta}} s}.$$

$Z_{mo}^{\text{sta,ml,b,g}}$ can be formulated as

$$Z_{mo}^{\text{sta,ml,b,g}} = (Z_{m+1,o}^{\text{sta,ml,b,ba}} + Z_{m+1,o}^{\text{sta,ml,b,g}\prime}) \exp(\beta^{\text{sta}} sp^{\text{og}}) + Z_{m+1,o}^{\text{sta,ml,b,g}} \exp(\beta^{\text{sta}} sp^{\text{eg}}).$$

$Z_{mo}^{\text{sta,ml,b,g}\prime}$ can be formulated as

$$Z_{mo}^{\text{sta,ml,b,og}\prime} = (Z_{m,o+1}^{\text{sta,ml,b,ba}} + Z_{m,o+1}^{\text{sta,ml,b,g}}) \exp(\beta^{\text{sta}} sp^{\text{og}}) + Z_{m,o+1}^{\text{sta,ml,b,g}\prime} \exp(\beta^{\text{sta}} sp^{\text{eg}})$$

Each of $Z_{jl}^{\text{sta,ml,b}}, Z_{jl}^{\text{sta,ml,b,ba}}, Z_{jl}^{\text{sta,ml,b,g}}, Z_{jl}^{\text{sta,ml,b,g}\prime}$ is formulated (or initialized) as each of

$$Z_{jl}^{\text{sta,ml,b}} = Z_{jl}^{\text{sta,ml,b,ba}} = Z_{jl}^{\text{sta,ml,b,g}} = Z_{jl}^{\text{sta,ml,b,g}\prime} = 0.$$

$Z_{mo}^{\text{sta,abp,b}}$ can be formulated as

$$Z_{mo}^{\text{sta,abp,b}} = Z_{mo}^{\text{sta,abp,b,ba}} + Z_{mo}^{\text{sta,abp,b,g}} + Z_{mo}^{\text{sta,abp,b,g}\prime}$$

where $Z_{mo}^{\text{sta,abp,b,ba}}$ is defined as $Z_{mo}^{\text{sta,abp,b}}$ in the case in which the two positions $m, o$ are aligned; $Z_{mo}^{\text{sta,abp,b,og}}$ $Z_{mo}^{\text{sta,abp,b}}$ in the case in which $m$ is aligned with a gap in front of $o$; and $Z_{mo}^{\text{sta,abp,b,og}\prime}$ $Z_{mo}^{\text{sta,abp,b}}$ in the case in which $o$ is aligned with a gap in front of $m$. $Z_{mo}^{\text{sta,abp,b,ba}}$ can be formulated as

$$Z_{mo}^{\text{sta,abp,b,ba}} = \sum_{np} (Z_{n+1,p+1}^{\text{sa,b}} \exp(-2\beta^{\text{sta}} fe^{\text{ml,abp}}) Z_{mnop}^{\text{sta,bpa}}) + Z_{m+1,o+1}^{\text{sta,abp,b}} (or_{mo}^{\text{ba}})^{\beta^{\text{sta}} s}.$$

$Z_{mo}^{\text{sta,abp,b,g}}$ can be formulated as

$$Z_{mo}^{\text{sta,abp,b,g}} = (Z_{m+1,o}^{\text{sta,abp,b,ba}} + Z_{m+1,o}^{\text{sta,abp,b,og}\prime}) \exp(\beta^{\text{sta}} sp^{\text{og}}) + Z_{m+1,o}^{\text{sta,abp,b,g}} \exp(\beta^{\text{sta}} sp^{\text{eg}}).$$

$Z_{mo}^{\text{sta,abp,b,g}\prime}$ can be formulated as

$$Z_{mo}^{\text{sta,abp,b,g}\prime} = (Z_{m,o+1}^{\text{sta,abp,b,ba}} + Z_{m,o+1}^{\text{sta,abp,b,g}}) \exp(\beta^{\text{sta}} sp^{\text{og}}) + Z_{m,o+1}^{\text{sta,abp,b,g}\prime} \exp(\beta^{\text{sta}} sp^{\text{eg}}).$$

Each of $Z_{jl}^{\text{sta,abp,b}}, Z_{jl}^{\text{sta,abp,b,ba}}, Z_{jl}^{\text{sta,abp,b,g}}, Z_{jl}^{\text{sta,abp,b,g}\prime}$ is formulated (or initialized) as each of

$$Z_{jl}^{\text{sta,abp,b}} = Z_{jl}^{\text{sta,abp,b,ba}} = Z_{jl}^{\text{sta,abp,b,g}} = Z_{jl}^{\text{sta,abp,b,g}\prime} = 0.$$

Lemma 2.1. *$Z^{\cdot}_{mo}$ can be computed on an inside algorithm with the time and space complexities $O(N^4M^4), O(N^2M^2)$.*

Proof. $Z^{\mathrm{sta,bpa}}_{ijkl}, Z^{\mathrm{sta,bpa},\lambda}_{ijkl}$ can be computed on an inside algorithm with the time and space complexities $O(N^4M^4), O(N^2M^2)$ from Supplementary lemma 1.1. $Z^{\cdot}_{mo}$ can be computed on a DP with at the most time and space complexities $O(NM), O(NM)$ since $Z^{\cdot}_{mo}$ involves at most positions $n, p$ and the two positions $m, o$ in the formulation of $Z^{\cdot}_{mo}$. Therefore, $Z^{\cdot}_{mo}$ can be computed on an inside algorithm with the time and space complexities $O(N^2M^2), O(N^2M^2)$.

#### 2.2 Formulation of base pair alignment probability $p^{\mathrm{bpa}}_{mnop|\mathbb{RNA}}$

$p^{\mathrm{bpa}}_{mnop|\mathbb{RNA}}$ can be formulated as

$$p^{\mathrm{bpa}}_{mnop|\mathbb{RNA}} = \begin{cases} 1 & (m = o = 0 \land n = N+1 \land p = M+1) \\ Z^{\mathrm{sta,bpa}}_{mnop} \sum_{ijkl} \{ \frac{p^{\mathrm{bpa}}_{ijkl|\mathbb{RNA}} (or^{\mathrm{bpa}}_{ijkl})^{\beta^{\mathrm{sta}} s}}{Z^{\mathrm{sta,bpa}}_{ijkl}} [Z^{\mathrm{sa,f}}_{m-1,o-1} \exp(-\beta^{\mathrm{sta}}(fe^{\mathrm{2l}}_{ijmn} + fe^{\mathrm{2l}}_{klop})) & \\ Z^{\mathrm{sa,b}}_{n+1,p+1} + \exp(-2\beta^{\mathrm{sta}}(fe^{\mathrm{ml,cbp}} + fe^{\mathrm{ml,abp}}))(Z^{\mathrm{sta,ml,f}}_{m-1,o-1} Z^{\mathrm{sa,b}}_{n+1,p+1} + Z^{\mathrm{sa,f}}_{m-1,o-1} & (otherwise) \\ Z^{\mathrm{sta,ml,b}}_{n+1,p+1} + Z^{\mathrm{sta,abp,f}}_{m-1,o-1} Z^{\mathrm{sa,b}}_{n+1,p+1} + Z^{\mathrm{sa,f}}_{m-1,o-1} Z^{\mathrm{sta,abp,b}}_{n+1,p+1} + Z^{\mathrm{sta,ml,f}}_{m-1,o-1} Z^{\mathrm{sta,abp,b}}_{n+1,p+1} & \\ + Z^{\mathrm{sta,abp,f}}_{m-1,o-1} Z^{\mathrm{sta,ml,b}}_{n+1,p+1} + Z^{\mathrm{sta,ml,f}}_{m-1,o-1} Z^{\mathrm{sta,ml,b}}_{n+1,p+1} + Z^{\mathrm{sta,abp,f}}_{m-1,o-1} Z^{\mathrm{sta,abp,b}}_{n+1,p+1})] \} & \end{cases} .$$

An interpretation of the formulation of $p^{\mathrm{bpa}}_{mnop|\mathbb{RNA}}$ is shown on (B) in Supplementary figure 3.

#### 2.3 Analysis of time and space complexities of computing base alignment probability matrix and base pair alignment probability matrix $\boldsymbol{P}^{\mathrm{ba,sta}}_{|\mathbb{RNA}}, \boldsymbol{P}^{\mathrm{bpa}}_{|\mathbb{RNA}}$

Theorem 2.1. *$\boldsymbol{P}^{\mathrm{ba,sta}}_{|\mathbb{RNA}}, \boldsymbol{P}^{\mathrm{bpa}}_{|\mathbb{RNA}}$ can be computed on a DP with the time and space complexities $O(N^4M^4), O(N^2M^2)$.*

Proof. $Z^{\mathrm{sta,bpa}}_{ijkl}, Z^{\mathrm{sta,bpa},\lambda}_{ijkl}$ can be computed on an inside algorithm with the time and space complexities $O(N^4M^4), O(N^2M^2)$ from Supplementary lemma 1.1. $Z^{\cdot}_{np}, Z^{\cdot}_{mo}$ can be computed on an inside algorithm with the time and space complexities $O(N^2M^2), O(N^2M^2)$ from Supplementary lemma 1.1 and Supplementary lemma 2.1. $p^{\mathrm{ba,sta}}_{np|\mathbb{RNA}}$ can be computed on a DP with the time and space complexities $O(N^2M^2), O(N^2M^2)$ since $p^{\mathrm{ba,sta}}_{np|\mathbb{RNA}}$ involves positions $i, j, k, l$ and the two positions $n, p$ in the formulation of $p^{\mathrm{ba,sta}}_{np|\mathbb{RNA}}$. $p^{\mathrm{bpa}}_{mnop|\mathbb{RNA}}$ can be computed on a DP with the time and space complexities $O(N^2M^2), O(N^2M^2)$ since $p^{\mathrm{bpa}}_{mnop|\mathbb{RNA}}$ involves positions $i, j, k, l$ and the four positions $m, n, o, p$ in the formulation of $p^{\mathrm{bpa}}_{mnop|\mathbb{RNA}}$. Therefore, $\boldsymbol{P}^{\mathrm{ba,sta}}_{|\mathbb{RNA}}, \boldsymbol{P}^{\mathrm{bpa}}_{|\mathbb{RNA}}$ can be computed on a DP with the time and space complexities $O(N^4M^4), O(N^2M^2)$.

DPs like this DP are called IO algorithms since recursions for these DPs are proceeded inside or outside.

### 3 Analysis of time and space complexities of Algorithm 1 with probable-STA restriction

Theorem 3.1. *$\boldsymbol{P}^{\mathrm{ba,sta,a}}_{|\mathbb{RNA}}, \boldsymbol{P}^{\mathrm{bpa,a}}_{|\mathbb{RNA}}$ can be computed with the time and space complexities $O(KJ(\delta^{\mathrm{m}})^2), O(K+J); K := L\delta\delta^{\mathrm{m}}, \delta := \delta^{\mathrm{bp}} g^{\mathrm{m}}, \delta^{\mathrm{bp}} := \lfloor \frac{1}{p^{\mathrm{bp,ss,m}}} \rfloor, \delta^{\mathrm{m}} := \min(\delta^{\mathrm{bp}}, g^{\mathrm{m}}), J := Lg^{\mathrm{m}}$ by Algorithm 1 with the probable-STA restriction.*

Proof. The number of $Z^{\mathrm{sta,bpa}}_{ijkl}, Z^{\mathrm{sta,bpa},\lambda}_{ijkl}$ satisfying the probable-STA restriction is at most $O(K)$. The number of $Z^{\cdot}_{np}, Z^{\cdot}_{mo}$ satisfying the maximum-gap-number is $O(J)$. Therefore, $\boldsymbol{P}^{\mathrm{ba,sta,a}}_{|\mathbb{RNA}}, \boldsymbol{P}^{\mathrm{bpa,a}}_{|\mathbb{RNA}}$ can be computed with the time and space complexities $O(KJ(\delta^{\mathrm{m}})^2), O(K+J)$ by Algorithm 1 with the probable-STA restriction.

### 4 Derivation of accuracy $a^{\gamma\mathrm{c}}_{\boldsymbol{SS}\mathbb{STA}}$

Corollary 4.1. *$\forall \boldsymbol{SS}\mathbb{STA}[2TP^{\mathrm{ss}} + FN^{\mathrm{ss}} = \sum_i I^{\mathrm{bp}}_{\mathbb{STA}i} \land TN^{\mathrm{ss}} + FP^{\mathrm{ss}} = \sum_i (1 - I^{\mathrm{bp}}_{\mathbb{STA}i})]$ is true.*

$a_{\boldsymbol{SS}\mathbb{STA}}$ can be formulated as

$$\begin{aligned} a_{\boldsymbol{SS}\mathbb{STA}} &= \alpha_1 TP^{\mathrm{ss}} + \alpha_2 TN^{\mathrm{ss}} - \alpha_3 [\sum_i (1 - I^{\mathrm{bp}}_{\mathbb{STA}i}) - TN^{\mathrm{ss}}] - \alpha_4 [\sum_i (I^{\mathrm{bp}}_{\mathbb{STA}i}) - 2TP^{\mathrm{ss}}] \\ &= c + (\alpha_1 + 2\alpha_4) TP^{\mathrm{ss}} + (\alpha_2 + \alpha_3) TN^{\mathrm{ss}} \\ &= c + (\alpha_2 + \alpha_3)(\gamma TP^{\mathrm{ss}} + TN^{\mathrm{ss}}) \\ &= c + c' a^{\gamma\mathrm{c}}_{\boldsymbol{SS}\mathbb{STA}} \\ c &:= -\alpha_3 \sum_i (1 - I^{\mathrm{bp}}_{\mathbb{STA}i}) - \alpha_4 \sum_i I^{\mathrm{bp}}_{\mathbb{STA}i}, c' := \alpha_2 + \alpha_3 \end{aligned} ; \quad (1)$$

with Supplementary corollary 4.1.

Corollary 4.2. *$a_{\boldsymbol{SS}\mathbb{STA}}, a^{\gamma\mathrm{c}}_{\boldsymbol{SS}\mathbb{STA}}$ are equivalent in terms of SS accuracy from Supplementary equation 1.*

#### 5 Derivation of expected accuracy $\mathbb{E}_{\mathtt{STA}}[a^{\gamma\mathrm{c}}_{\boldsymbol{SS}\mathtt{STA}}]$

$\mathbb{E}_{\mathtt{STA}}[a^{\gamma\mathrm{c}}_{\boldsymbol{SS}\mathtt{STA}}]$ can be formulated as

$$\begin{aligned}
\mathbb{E}_{\mathtt{STA}}[a^{\gamma\mathrm{c}}_{\boldsymbol{SS}\mathtt{STA}}] &:= \sum_{\mathtt{STA}} (a^{\gamma\mathrm{c}}_{\boldsymbol{SS}\mathtt{STA}} p_{\mathtt{STA}}) \\
&= \sum_{\mathtt{STA}} (\{\gamma \sum_{ij} (I^{\mathrm{bp}}_{\boldsymbol{SS}ij} I^{\mathrm{bp}}_{\mathtt{STA}ij}) + \sum_{i} [(1 - I^{\mathrm{bp}}_{\boldsymbol{SS}i})(1 - I^{\mathrm{bp}}_{\mathtt{STA}i})]\} p_{\mathtt{STA}}) \\
&= \sum_{\mathtt{STA}} \{[\gamma \sum_{ij \mid I^{\mathrm{bp}}_{\boldsymbol{SS}ij}=1} (I^{\mathrm{bp}}_{\mathtt{STA}ij}) + \sum_{i \mid I^{\mathrm{bp}}_{\boldsymbol{SS}i}=0} (1 - I^{\mathrm{bp}}_{\mathtt{STA}i})] p_{\mathtt{STA}}\} \\
&= \gamma \sum_{ij \mid I^{\mathrm{bp}}_{\boldsymbol{SS}ij}=1} ( \sum_{\mathtt{STA} \mid I^{\mathrm{bp}}_{\mathtt{STA}ij}=1} p_{\mathtt{STA}}) + \sum_{i \mid I^{\mathrm{bp}}_{\boldsymbol{SS}i}=0} \sum_{\mathtt{STA} \mid I^{\mathrm{bp}}_{\mathtt{STA}i}=0} p_{\mathtt{STA}} \\
&= \gamma \sum_{ij \mid I^{\mathrm{bp}}_{\boldsymbol{SS}ij}=1} (p^{\mathrm{bp,sta}}_{ij \mid \mathbb{RNA}}) + \sum_{i \mid I^{\mathrm{bp}}_{\boldsymbol{SS}i}=0} p^{\mathrm{up,sta}}_{i \mid \mathbb{RNA}}.
\end{aligned}$$

#### 6 Algorithms for computing maximum-expected-accuracy secondary structure $\widehat{\boldsymbol{SS}}_{\boldsymbol{RNA}}$

Lemma 6.1. *$mea_{ij}$ can be computed on a DP with the time and space complexities $O(N^4), O(N^2)$.*

Proof. $mea_{ij}$ can be computed on a DP with the time and space complexities $O(1), O(1)$ since $mea_{ij}$ involves only the two indexes $i, j$ in Equation 4. $mea_n$ can be computed on a DP with the time and space complexities $O(N), O(N)$ since $mea_n$ involves indexes and the index $m, n$ in Equation 4. $mea_n$ can be discarded after $mea_{ij}$ is computed according to Equation 4. Therefore, $mea_{ij}$ can be computed on a DP with the time and space complexities $O(N^4), O(N^2)$.

Lemma 6.2. *$\widehat{\boldsymbol{SS}}_{\boldsymbol{RNA}}$ can be computed from the MEA matrix $\boldsymbol{MEA} := (mea_{ij})$ with the time and space complexities $O(N^4), O(N^2)$ by Supplementary algorithm 1.*

**Algorithm 1** A traceback algorithm for computing a maximum-expected-accuracy secondary structure $\widehat{\boldsymbol{SS}}_{\boldsymbol{RNA}}$.

**function** tracebackForMeaSs($\boldsymbol{MEA}$)
  Initialize MEA SS components $\widehat{ss}_{ij}$; $(\widehat{ss}_{ij}) := \widehat{\boldsymbol{SS}}_{\boldsymbol{RNA}}$ as 0s
  Set $\widehat{ss}_{0,N+1}$ to 1
  Prepare an empty stack $\boldsymbol{S}$
  Push the pseudo-position pair $[0, N+1]$ to $\boldsymbol{S}$
  **while** $\boldsymbol{S}$ is not empty **do**
    Pop a position pair $[i, j]$ from $\boldsymbol{S}$
    **if** $mea_{ij} = 0$ **then**
      **continue**
    **else**
      Compute $mea_n$ according to Equation 4
      $n \leftarrow j - 1$
      **while** $mea_n > 0$ **do**
        **if** $mea_n = mea_{n-1} + p^{\mathrm{up,sta}}_{n \mid \mathbb{RNA}}$ **then**
          $n \leftarrow n - 1$
        **else**
          **for** m **do**
            **if** $mea_n = mea_{m-1} + mea_{mn}$ **then**
              Set $\widehat{ss}_{mn}$ to 1
              Push the position pair $[m, n]$ to $\boldsymbol{S}$
              $n \leftarrow m - 1$
              **break**
  **return** $\widehat{\boldsymbol{SS}}_{\boldsymbol{RNA}}$

Proof. $t(\boldsymbol{S})$ is $O(N^2)$ since $\boldsymbol{S}$ involves indexes $i, j$. The operations in each step of the iteration for $\boldsymbol{S}$ require the time and space complexities $O(N^2), O(N^2)$ since $mea_n$ can be computed with the time and space complexities $O(N^2), O(N^2)$ and the iteration for $mea_n$ involves indexes and the index $m, n$. Therefore, $\widehat{\boldsymbol{SS}}_{\boldsymbol{RNA}}$ can be computed from $\boldsymbol{MEA}$ with the time and space complexities $O(N^4), O(N^2)$ by Supplementary algorithm 1.

Corollary 6.1. *$\widehat{\boldsymbol{SS}}_{\boldsymbol{RNA}}$ can be computed with the time and space complexities $O(N^4), O(N^2)$ by Supplementary algorithm 2.*

Corollary 6.2. *$\widehat{\boldsymbol{SS}}^{\mathrm{pct}}_{\boldsymbol{RNA}}$ can be computed by the NeoFold algorithm, Supplementary algorithm 3, with the time and space complexities $O(H^2), O(H)$.*

**Algorithm 2** An algorithm for computing a maximum-expected-accuracy secondary structure $\widehat{\boldsymbol{SS}}_{\boldsymbol{RNA}}$.

**function** getMeaSs($(p^{\mathrm{bp,sta}}_{ij|\mathbb{RNA}}), (p^{\mathrm{up,sta}}_{i|\mathbb{RNA}})$)
  Compute $mea_{ij}, mea_n$ on a DP according to Equation 4
  **return** tracebackForMeaSs($\boldsymbol{MEA}$)

**Algorithm 3** The NeoFold algorithm.

**function** neoFold($\boldsymbol{P}^{\theta',\mathrm{pct}}_{|\boldsymbol{RNA},\boldsymbol{RF}}$)
  **return** getMeaSs($\boldsymbol{P}^{\theta',\mathrm{pct}}_{|\boldsymbol{RNA},\boldsymbol{RF}}$)

#### 7 Principle of each of CentroidHomFold and TurboFold-MEA algorithms

The CentroidHomFold algorithm maximizes EA

$$\sum_{ij|I^{\mathrm{bp}}_{\boldsymbol{SS}ij}=1} [(\gamma^{\mathrm{ch}} + 1)p^{\mathrm{bp,pct,d}}_{ij|\boldsymbol{RNA}} - 1]; \gamma^{\mathrm{ch}} \in \Re^+, p^{\mathrm{bp,pct,d}}_{ij|\boldsymbol{RNA}} := \frac{p^{\mathrm{bp,ss}}_{ij|\boldsymbol{RNA}} + (R-1)\sum_{\boldsymbol{RNA'} \in \boldsymbol{RF} \setminus (\boldsymbol{RNA}), kl} (p^{\mathrm{ba,sa}}_{ik|\mathbb{RNA}} p^{\mathrm{ba,sa}}_{jl|\mathbb{RNA}} p^{\mathrm{bp,ss}}_{kl|\boldsymbol{RNA'}})}{R}.$$

The TurboFold-MEA algorithm maximizes EA

$$2\gamma^{\mathrm{t}} \sum_{ij|I^{\mathrm{bp}}_{\boldsymbol{SS}ij}=1} (p^{\mathrm{bp,ss,t}}_{ij|\boldsymbol{RNA}}) + \sum_{i|I^{\mathrm{bp}}_{\boldsymbol{SS}i}=0} p^{\mathrm{up,t}}_{i|\boldsymbol{RNA}}; \gamma^{\mathrm{t}} \in \Re^+$$

where $p^{\mathrm{bp,ss,t}}_{ij|\boldsymbol{RNA}}$ is defined as the BPP of the two positions $i, j$ given $\boldsymbol{RNA}$ on SS refined iteratively on this algorithm and $p^{\mathrm{up,t}}_{i|\boldsymbol{RNA}}$ the unpairing probability of $i$ given $\boldsymbol{RNA}$ on SS refined iteratively on this algorithm. This algorithm utilizes $p^{\mathrm{bp,ss,t}}_{ij|\boldsymbol{RNA}}, p^{\mathrm{up,t}}_{i|\boldsymbol{RNA}}$ under this refinement for computing ML pairwise SAs. This algorithm utilizes these SAs for refining $p^{\mathrm{bp,ss,t}}_{ij|\boldsymbol{RNA}}, p^{\mathrm{up,t}}_{i|\boldsymbol{RNA}}$ more under this refinement. These two utilizations are iterated until $p^{\mathrm{bp,ss,t}}_{ij|\boldsymbol{RNA}}, p^{\mathrm{up,ss,t}}_{i|\boldsymbol{RNA}}$ are refined fully.

#### 8 Tables and figures

Table 1. The seven parameter values of the RNAfamProb program.

| Parameter | Value |
|---|---|
| Scale parameter $\beta^{\mathrm{sta}}$ | $\frac{1}{RT}$ $[kcal/mol]; R := 1.98717 \cdot 10^{-3}$ $[kcal/(K \cdot mol)], T := 310.15$ $[K]$ |
| Scale parameter $s$ | $\frac{1}{\beta^{\mathrm{sta}}}$ $[mol/kcal]$ |
| Minimum BPP $p^{\mathrm{bp,ss,m}}$ | 0.005 |
| Maximum gap-number in STA $g^{\mathrm{m}}$ | $\lvert N - M \rvert$ |
| Opening gap penalty $p^{\mathrm{og}}$ | 0 |
| Extending gap penalty $p^{\mathrm{eg}}$ | 0 |
| Number of threads | Number of available threads |

Table 2. The number of the RNA sequences and range of the RNA sequence lengths on each ncRNA family in test set 1.

| NcRNA family | Number of RNA sequences | Range of RNA sequence lengths [nt] |
|---|---|---|
| RNAIII | 4 | 489–517 |
| Hammerhead ribozyme (type III) | 10 | 40–58 |
| Hammerhead ribozyme (type I) | 10 | 45–118 |
| Bicoid 3 prime-UTR regulartory element | 4 | 543–551 |
| R2 RNA element | 10 | 202–237 |
| Gammaretrovirus core encapsidation signal | 10 | 100–102 |
| RNase MRP | 5 | 246–305 |
| Ciliate telomerase RNA | 10 | 160–215 |
| Coronavirus 3 prime stem-loop II-like motif (s2m) | 10 | 43 |
| Vimentin 3 prime UTR protein-binding region | 10 | 65–77 |
| RNase E 5 prime UTR element | 6 | 337–339 |

```
(A)
GCCCGGGUGGUGAAAUCGGUAGACACGCAGGACUUAAAAUCCUGUGGCAUAAAAGCCAUGUCGGUUCAAGUCCGACCCCGGGCA
(((((((..(((...((....)))))((((((.......))))))(((......)))..(((((.......))))))))))))).
GCCGAAAUAGCUCAGUUGGGAGAGCGUUAGACUGAAGAUCUAAAAGGUCCCUGGUUCGAUCCGGGUUUCGGCA
(((((((....((........)).............(((((...)))))(((((.......)))))))))))).
GACUCGUUAGCUCAGCCGGUAGAGCAACUGGCUUUUAACCAGUGGGUCCGGGGUUCGAAUCCCCGACGAGUCA
(((((((......(((((((......)))))))....(((....))).(((((.......)))))))))))).
GGCUUUUUAGCUCAGCAGGUAGAGCAACCGGCUGUUAACCGGUUUGUCACAGGUUCGAGCCCUGUAAAAGCCG
(((((((..(((((((..((.((..((((((.......))))))..))))..))).)))).....))))))).
UCUUUCUAGUACUAAGGAGUAUAAGUGGCUUCCAACCACACGGUCUUGGUUAGAGUCCAAGGAAAGAU
(((((((.(((((....)))))...(((.(((.(((((........))))).))).))).))))))).
GGAGGAAUACCCAAGUCUGGCUGAAGGGAUCGGUCUUGAAAACCGACAGGGUGUCAAAGCCCGCGGGGGUUCGAAUCCCUCUUCCUCCG
(((((((..(((.............))).(((((.......)))))..((((......))))....(((.......)))..))))))).
(B)
GCCCGGGUGGUGAAAUCGGUAGACACGCAGGACUUAAAAUCCUGUGGCAUAAAAGCCAUGUCGGUUCAAGUCCGACCCCGGGCA
(((((((..(((...((....)))))((((((.......))))))(((......)))..(((((.......))))))))))))).
GCCGAAAUAGCUCAGUUGGGAGAGCGUUAGACUGAAGAUCUAAAAGGUCCCUGGUUCGAUCCGGGUUUCGGCA
(((((((....((........)).............(((((...)))))(((((.......)))))))))))).
GACUCGUUAGCUCAGCCGGUAGAGCAACUGGCUUUUAACCAGUGGGUCCGGGGUUCGAAUCCCCGACGAGUCA
(((((((......(((((((......)))))))....(((....))).(((((.......)))))))))))).
GGCUUUUUAGCUCAGCAGGUAGAGCAACCGGCUGUUAACCGGUUUGUCACAGGUUCGAGCCCUGUAAAAGCCG
(((((((..(((((((..((.((..((((((.......))))))..))))..))).)))).....))))))).
UCUUUCUAGUACUAAGGAGUAUAAGUGGCUUCCAACCACACGGUCUUGGUUAGAGUCCAAGGAAAGAU
(((((((.(((((....)))))...(((.(((.(((((........))))).))).))).))))))).
GGAGGAAUACCCAAGUCUGGCUGAAGGGAUCGGUCUUGAAAACCGACAGGGUGUCAAAGCCCGCGGGGGUUCGAAUCCCUCUUCCUCCG
(((((((..(((.............))).(((((.......)))))..((((......))))....(((.......)))..))))))).
p >= 5.00e-01, p >= 2.50e-01, p >= 1.25e-01, p >= 6.25e-02, p < 6.25e-02 where p is a
 posterior (pseudo-)base-pairing-probability for a base pair
```

**Fig. 4.** Six examples of secondary structures computed by the RNAfold program, an maximum-likelihood secondary-structure program (Lorenz et al., 2011), color-coded based on base-pairing probabilities $p^{\mathrm{bp,ss}}_{ij|\boldsymbol{RNA}}$ and pseudo-base-pairing-probabilities $p^{\mathrm{bp,pct}}_{ij|\boldsymbol{RNA},\boldsymbol{RF}}$. (A) These six secondary structures color-coded based on base-pairing probabilities $p^{\mathrm{bp,ss}}_{ij|\boldsymbol{RNA}}$ computed by the McCaskill program. (B) These six secondary structures color-coded based on pseudo-base-pairing-probabilities $p^{\mathrm{bp,pct}}_{ij|\boldsymbol{RNA},\boldsymbol{RF}}$ computed by the RNAfamProb suite. A “()” means the two positions corresponding to these two parentheses are BP. These six sequences are the same as Figure 5. The RNAfamProb program was run with the seven parameter values same as Figure 5.
